# Single-cell transcriptome analysis of the immunosuppressive effect of differential expression of tumor PD-L1 on responding TCR-T cells

**DOI:** 10.1101/2020.07.23.217059

**Authors:** Renpeng Ding, Shang Liu, Shanshan Wang, Huanyi Chen, Fei Wang, Qumiao Xu, Linnan Zhu, Xuan Dong, Ying Gu, Cheng-Chi Chao, Qianqian Gao

## Abstract

PD-L1 expression levels in tumors do not consistently predict cancer patients’ response to PD-(L)1 inhibitors. We therefore evaluated how tumor PD-L1 levels affect the anti-PD-(L)1 efficacy and T cell function. We used MART-1-specific TCR-T cells (TCR-T_MART-1_) stimulated with MART-1_27-35_ peptide-loaded MEL-526 tumor cells with different proportions of them expressing PD-L1 to perform cellular assays and high-throughput single-cell RNA sequencing. Compared to control T cells, TCR-T_MART-1_ were more sensitive to exhaustion and secreted lower pro-inflammatory but higher anti-inflammatory cytokines with increasing proportions of PD-L1^+^ tumor cells. The colocalization of T cells and tumor cells in gene clusters correlated negatively with the proportion of PD-L1^+^ tumor cells and positively with immune cell cytotoxicity. Moreover, elevated proportion of PD-L1^+^ tumor cells increased PD-L1 expression and decreased PD-1 expression on T cells and enhanced T cell death. The expression of PD-1 and PD-L1 in T cells and macrophages also correlated positively with COVID-19 severity.

## Introduction

Programmed cell death-ligand-1 (PD-L1) is the ligand of programmed death-1 (PD-1), which are encoded by *CD274* and *PDCD1*, respectively. PD-L1 is expressed in many cancer tissues, including melanoma [1], a widely recognized immunogenic neoplasm. Expression of PD-L1 is undetectable in most normal tissues, but can be induced by inflammatory cytokines, especially interferon-γ (IFN-γ) in various cell types [2-4]. As a strategy to evade immune responses, PD-L1 is often up-regulated on tumor cells and induces T cell anergy, exhaustion or apoptosis upon engagement with PD-1 expressed on tumor infiltrating lymphocytes (TILs) to impair T cell responses[1, 5]. Expression of PD-L1 is not restricted to tumor cells, PD-L1 is also expressed in TILs and its expression by TILs correlates with aggressive tumors, demonstrating the immunosuppressive role of PD-L1 [6, 7]. Binding of PD-1 and PD-L1 impairs T cell activation by interfering with Ras-Raf-MEK-ERK and PI3K-AKT signaling pathways that promote T cell proliferation and differentiation [8]. In addition to binding PD-1, PD-L1 has been reported to interact with CD80 in *cis* to modulate T cell function and tumor microenvironment [9, 10].

The PD-1/PD-L1 signaling pathway plays an important role in tumor evasion from host immune responses [11]. Inhibitors of PD-1 and PD-L1 have been studied in various tumor types and have now been approved for treating many malignancies, including melanoma, non–small-cell lung cancer (NSCLC), and bladder cancer. [12-16] PD-L1 expression on tumor cells and tumor infiltrating antigen presenting cells (APCs) has been approved as a companion biomarker for the treatment with some of these inhibitors [17-22]. Positive correlation between higher level of PD-L1 expression and higher response rate in melanoma has also been demonstrated [23-25]. However, some studies showed that PD-L1 expression is insufficient to predict a benefit from immune checkpoint inhibitor (CPI) therapy and PD-L1 expression level alone is a poor predictive biomarker of overall survival [26, 27].

The PD-L1 expression level has different predictive values for response to PD-(L)1 blockade in different types of tumors, many tumors that express PD-L1 do not respond to PD-1 or PD-L1 inhibitors. The overall low response rates of PD-1 and PD-L1 inhibitors limit their clinical application. Thus, it is important to know how PD-L1 and its expression level on tumor cells affect the efficacy of immunotherapy and T cell function. The role of PD-L1 has been studied for many years [4-6, 22, 28], but only from the bulk T cell level, which is hard to elucidate the exact relationship between PD-L1 expression and T cell function.

In this study, we used high-throughput single-cell mRNA sequencing (scRNA-seq), multiplex cytokine secretion assay, and cell cytotoxicity assays to investigate the immunoregulatory effect of tumor PD-L1 on responding TCR-T cells. Our research is the first to dissect at the single-cell level transcriptional features as well as cytokine and cytotoxic signatures of antigen-specific TCR-T cells responding to different tumor PD-L1 ratios. Furthermore, single-cell immune profiling was explored in COVID-19 patients, which is essential for understanding the potential mechanisms underlying COVID-19 pathogenesis.

## Results

### Increased tumor PD-L1 expression suppressed cytotoxicity and cytokine secretion of TCR-T_MART-1_

We used cytotoxicity and cytokine secretion assays together with scRNA-seq to interrogate TCR-T cells stimulated by MEL-526 melanoma cells with different proportions of them expressing PD-L1 (Fig. 1A). This approach made it possible to quantitatively dissect the T-cell activation state in relation to their subtypes, gene expression and cell differentiation. HLA-A*0201/Melan-A-specific TCR sequence (designed as TCR_MART-1_) was attained from T cells stimulated with Melan-A (aa27-35, LAGIGILTV) peptide (data unpublished). Melan-A, also known as MART-1, is a melanocytic marker [29]. Human TCRα and TCRβ sequences fused with murine TCR constant region were synthesized and cloned into a lentiviral vector (Fig. S1A). T cells that expressed or did not express TCR_MART-1_ are designated as TCR-T_MART-1_ and T_null_, respectively. T_total_ represents the entire T cell population that includes both TCR-T_MART-1_ and T_null_. After lentiviral transduction into CD8^+^ T cells, 17.5% of T_total_ was TCR-T_MART-1_, which reached 97.2% after fluorescence-activated cell sorting (FACS) (Fig. S1B). To verify the cytolytic capacity, TCR-T_MART-1_ were stimulated by peptide-loaded MEL-526 cells or a mock control at the effector:target (E:T) ratio of 1:1. Compared to T_null_, TCR-T_MART-1_ killed MEL-526 cells efficiently when MEL-526 cells were loaded with the MART-1_27-35_ peptide (Fig. 1B). TCR-T_MART-1_ similarly killed T2 cells, another target cell line (Fig. S1C).

**Fig 1.**
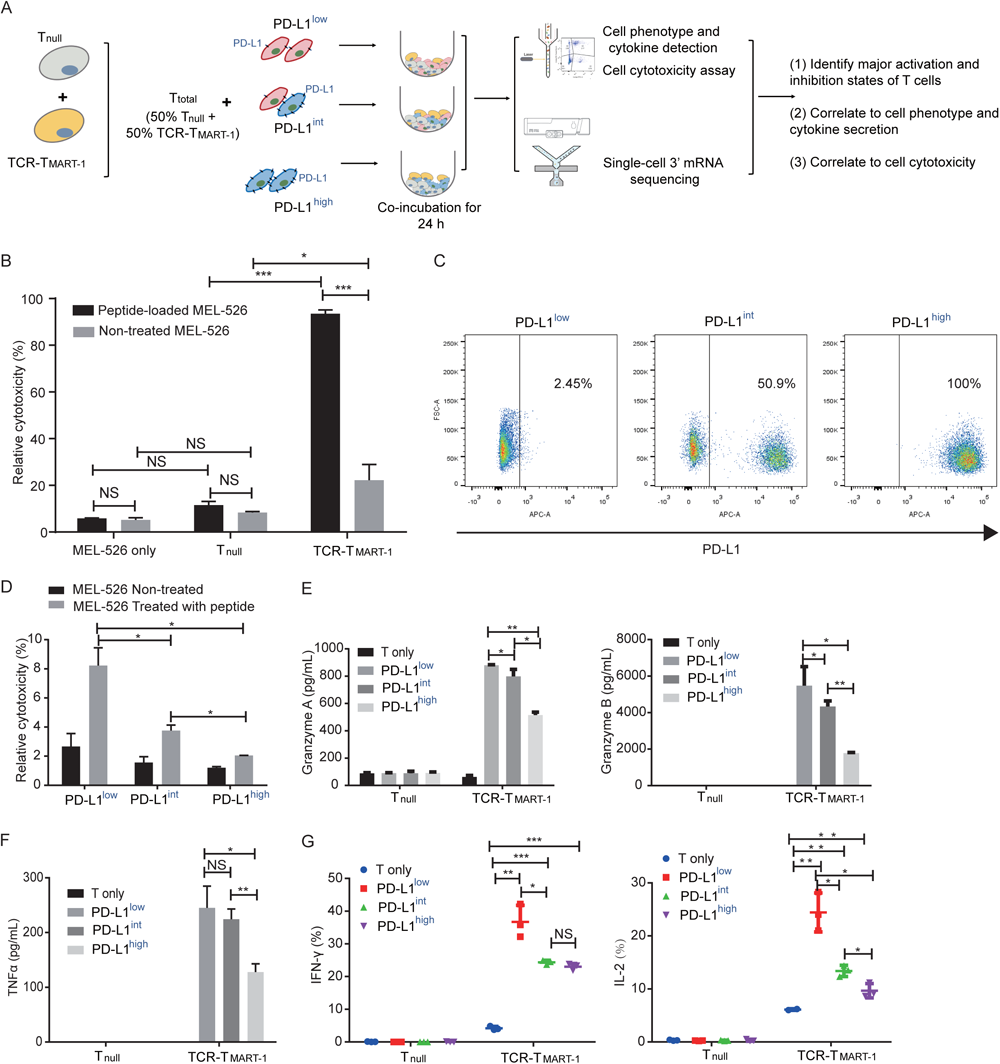
PD-L1 expression on melanoma MEL-526 cells pulsed with MART-1_26-35_ peptide inhibited cytotoxicity and cytokine secretion of TCR-T_MART-1_. (**A)** Overview of the study design. T_null_, control T cells; TCR-T_MART-1_, MART-1 specific TCR-T cells; T_total_, includes both T_null_ and TCR-T_MART-1_.**(B)** TCR-T_MART-1_ cytotoxicity against MEL-526 cells loaded with MART-1_26-35_ peptide or not at E:T ratio of 1:1. Error bars represent S.E.M. (*) 0.01<P < 0.05, (**) 0.001<P < 0.01, (***) P < 0.001. NS, not significant. **(C)** Flow cytometric analysis of PD-L1 expression on PD-L1^low^-, PD-L1^int^- and PD-L1^high^ MEL-526 cells. **(D)** TCR-T_MART-1_ cytotoxicity was inhibited by tumor PD-L1 in a dose dependent manner. T and TCR-T cells were incubated with different proportions of PD-L1^+^ MEL-526 cells for 24 h. **(E)** Secretion of Granzme A and Granzyme B by TCR-T_MART-1_ was inhibited by increased tumor PD-L1. T_null_ and TCR-T_MART-1_ were co-cultured with MART-1_26-35_ peptide loaded-MEL526 cells with different proportions of PD-L1 expression at E:T ratio of 1:1, and the secretion was detected by Cytometric Bead Array (CBA) system. **(F)** Secretion of TNF-α by TCR-T_MART-1_ was inhibited by increased proportion of PD-L1 expression among MEL-526 cells. **(G)** Secretion of IFN-γ and IL-2 by TCR-T_MART-1_ was inhibited by increased percentage of PD-L1 expression among MEL-526 cells.

To investigate the immunosuppressive role of tumor PD-L1, PD-L1 was overexpressed (OE) on MEL-526 cells (Fig. S1D). Different percentages of PD-L1 positive tumor cells were obtained by mixing OE with wild-type (WT) MEL-526 cells based on the clinical PD-L1 expression ratio [30]. Three tumor cell populations with different percentages of MEL-526 expressing PD-L1 were used in the study: PD-L1^low^ (without exogenous PD-L1, 2.45%), PD-L1^int^ (intermediate, 50.9%), and PD-L1^high^ (high, 100%) (Fig. 1C). The cytolytic activity of TCR-T_MART-1_ was inhibited by increasing percentages of tumor cells expressing PD-L1 (Fig. 1D), demonstrating a dose-dependent suppression of PD-L1 on TCR-T cell cytotoxicity. PD-L1 also dose-dependently suppressed the secretion of Granzymes (Fig. 1E) and pro-inflammatory cytokines, including TNF α (Fig. 1F), IFNγ and IL2 (Fig. 1G), in T_null_ and TCR-T_MART-1_. Altogether, PD-L1-mediated immune suppression modulated cytotoxicity and cytokine secretion of MART-1-specific TCR-T cells.

### Single-cell level analysis of T cells responding to peptide-pulsed MEL-526 cells

Single-cell transcriptome profiling was performed using a negative pressure orchestrated DNBelab C4 system [31]. Transcriptome profiling of a total of 20888 cells from four conditions was obtained after filtering out cells with low quality (Fig. 2A). To investigate the intrinsic T cell heterogeneity, unsupervised clustering was performed (Fig. 2B). T and tumor cells were identified by the expression of classic cell type markers, including *PTPRC, CD3D, CD3G, CD3E, CD8A, CD8B, TRAC, TRBC1*, and *TRBC2* for T cells and *MAGEA 4* for MEL-526 cells (Fig. 2C). Based on the expression of signature genes, T cells were composed of clusters 1, 3, 5, 6, 7, 9, 10, 11, 12, 14, 15, 16, and 19 (Fig. 2B). Exogenous TCR_MART-1_ was detected in cluster 6, 7, 11, 15, 16, and 19, but very little in cluster 10 and 12 (Fig. S1E). Furthermore, differentially expressed genes (DEGs) and known functional markers indicated the clusters of naïve, proliferating, early activated, cytotoxic, and exhausted CD8^+^ T cells (Fig. 2B).

**Fig 2.**
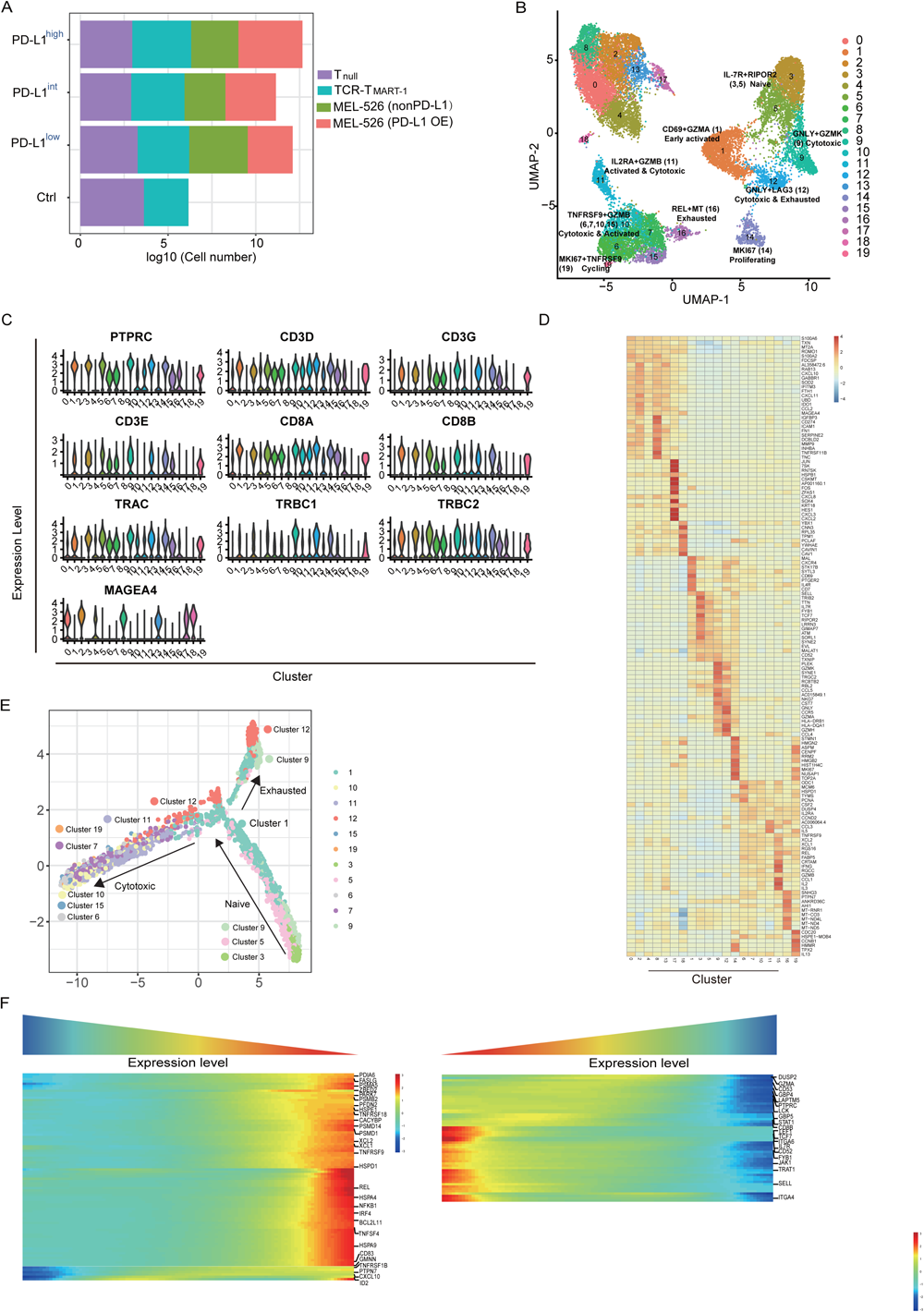
Single-cell level analysis of T cells responding to peptide-pulsed MEL-526 cells. **(A)** Cell number of T_null_, TCR-T_MART-1_, MEL-526 (non PD-L1), and MEL-526 (PD-L1 OE). **(B)** The UMAP projection of T cells and tumor cells, showing 20 main clusters in different colors. The phenotype description of each cluster is determined by marker gene expression of T cells and tumor cells. **(C)** Violin plots showing the expression profile of marker genes of T cells and tumor cells in the 20 clusters. **(D)** Heatmap of the 20 clusters with unique signature genes. **(E)** The ordering of T cells along pseudotime in a two-dimensional state-space defined by Monocle2. Cell orders were inferred from the expression of most dispersed genes across T cell populations. Each point corresponds to a single cell, and each color represents a T cell cluster. **(F)** The expression of genes was changed along the cell order.

DEG analysis further identified tumor cell clusters 0, 2, 4, 8, 13, 17, and 18 that showed high expressions of *S100A6, MAGEA 4*, and *HSPB1* as well as chemokines such as *CXCL10* and *CXCL11* (Fig. 2D). CXCL10 and CXCL11 recruit T cells and promote antitumor activity [32, 33]. For T cell clusters, *CXCR4* and early activation marker *CD69* were upregulated in cluster 1. Cluster 3 was more similar to cluster 5 and the expression of *SELL, IL7R*, and *TCF7* was upregulated, indicating a naïve phenotype. Expression of classic cytotoxic genes including *GZMK, NKG7, CST7, GNLY*, and *GZMA* were increased in cluster 9 and 12 while the expression of cell proliferation gene *MKI67* was upregulated in cluster 14. Expression of interleukins *IL5, IL2*, and *IL3* as well as of T cell activation and cytotoxicity genes such as *CSF2, XCL2, XCL1, IFNG*, and *GZMB* were upregulated in the remaining T cell clusters (Fig. 2D). According to the above characteristics (Fig. 2D), clusters shared similarities with each other were grouped together (Fig. 2B) and their DEGs were showed in Fig. S2A.

To understand T cell state transitions, an unsupervised inference method Monocle 2 [34] was applied to construct the potential development trajectories of ten T cell clusters (cluster 14, 19, and 16 were excluded due to their distinct expression of *MKI67* or mitochondrial genes). Cells from all clusters aggregated according to expression similarities to form a relative process in pseudotime, which began with cluster 3 and 5 (IL7R+RIPOR2, naïve cells), followed by cluster 9 (GNLY+GZMK) and 1 (CD69+GZMA) (Fig. 2E). Cluster 6, 7, 10, 15 (TNFRSF9+GZMB) and 19 (MKI67+TNFRSF9) activated and cytotoxic cells were located in the opposite directions with cluster 12 (GNLY+LAG3) in the pseudotime trajectory plot, demonstrating diverse functions of these cells. According to the trajectory analysis, CD8^+^ exhausted T cells were more closely linked to intermediate populations cluster 1 and 9 marked by GZMA and GZMK signatures, respectively than to the effector populations (Fig. 2E), consistent with a previous study [35]. Moreover, two main categories of genes were identified in this pseudotime process, one increased and the other one decreased (Fig. 2F). Upregulated genes included T cell activation and cytotoxicity markers, such as *XCL2, XCL1, TNFRSF9*, and *NFKB1*, and cell death associated genes, such as *FASLG* and *BCL2L11*. Downregulated genes encompassed naïve marker genes such as *LEF1, TCF7, IL7R*, and *SELL* (Fig. 2F). Taking together, while T cells differentiated into cytotoxic and exhausted populations, the expression of genes related with T cell activation and cytotoxicity was upregulated and the expression of naïve marker genes was downregulated in this pseudotime axis.

### Tumor PD-L1 affected T_null_ and TCR-T_MART-1_ differently at the transcriptional level

To reveal the structure of the overall T cell population, T cells were divided into T_null_ and TCR-T_MART-1_ and their cluster compositions were investigated. Cluster composition of the control (Ctrl) group was different from that of groups stimulated by tumor cells (Fig. 3A). After stimulation, cluster 1 (CD69+GZMA), 6&7&10&15 (TNFRSF9+GZMB), 11 (IL2RA+GZMB), 12 (GNLY+LAG3), and 16 (REL+MT) were increased compared to those of the Ctrl group (Fig. 3A), indicating the percentage of cytotoxic and exhausted T cells were increased upon antigen stimulation. With the increased percentage of tumor cells expressing PD-L1, only cluster 11 was increased in T_null_, whereas clusters 1, 11, and 12 were increased and clusters 6&7&10&15 were decreased in TCR-T_MART-1_. (Fig. 3A). These results implied that TCR-T_MART-1_ were more sensitive than T_null_ to increasing levels of tumor PD-L1, which also reduced the percentage of activated and cytotoxic TCR-T_MART-1_.

**Figure 3.**
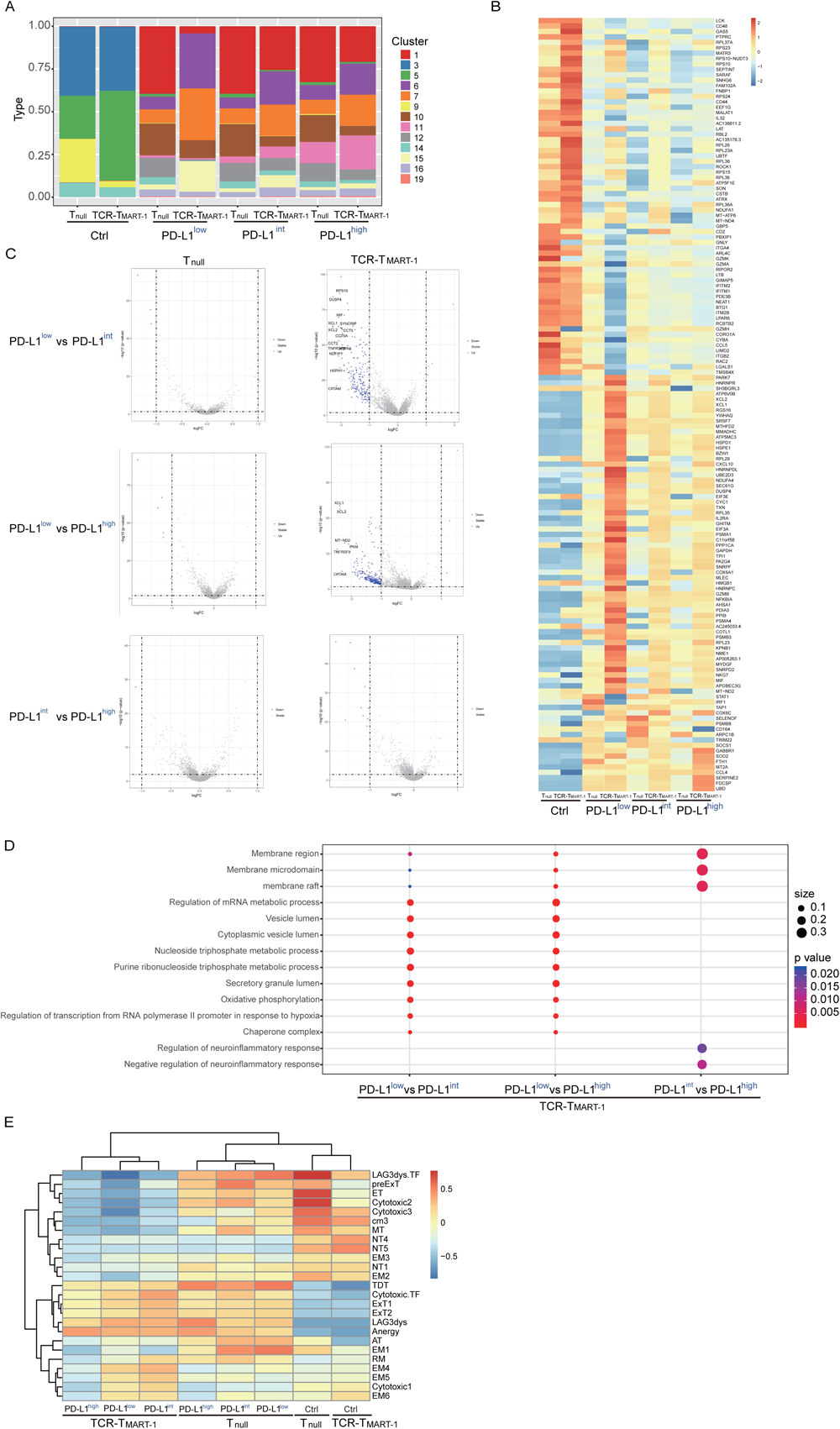
Cluster composition and gene expression analysis of T cells responding to different levels of tumor PD-L1. **(A)** Cluster composition of T_null_ and TCR-T_MART-1_. **(B)** Heatmap showing T_null_ and TCR-T_MART-1_ with unique signature genes. **(C)** Volcano plot showing differentially expressed genes in T_null_ (left) and TCR-T_MART-1_ (right) responding to differential proportion of PD-L1^+^ tumor. The cutoff is |logFC| >= 1 and p.value < 0.01. **(D)** Bubble plot showing the top 10 pathways in T_null_ (left) and TCR-T_MART-1_ (right) compared to the control group, respectively. The color represents pvalue and the size represents gene ratio. (**E)** GSVA analysis of cell differentiation status of T_null_ and TCR-T_MART-1_.

Results of gene expression (Fig. 3B, S3A) were consistent with the result of cluster composition in each group (Fig. 3A). TCR-T_MART-1_ were affected more than T_null_ by increased tumor PD-L1 (Fig. 3B, S3A, 3C), which inhibited the expression of T cell activation and cytotoxicity genes, including *XCL2, XCL1, IL2RA, GZMB*, and *NKG7* in TCR-T_MART-1_ (Fig. 3B). Moreover, there was no significant difference in gene expression between TCR-T_MART-1_ targeting PD-L1^int^ and PD-L1^high^ (Fig. 3C). Enriched signaling pathways were then analyzed. Different signaling were enriched in T_null_ and TCR-T_MART-1_ after encountering tumor cells (Fig. S3B). Compared to TCR-T_MART-1_ targeting PD-L1^int^ and PD-L1^high^, TCR-T_MART-1_ targeting PD-L1^low^ enriched metabolic and vesicle lumen related signaling (Fig. 3D). However, distinct pathways including membrane region, membrane microdomain and raft were enriched in TCR-T_MART-1_ targeting PD-L1^int^ compared to that targeting PD-L1^high^ (Fig. 3D). In addition, gene set variation analysis (GSVA) revealed that functional subtypes of T_null_ and TCR-T_MART-1_ populations responded to tumor cells differently. T_null_ were enriched with cytotoxic and terminally differentiated cells whereas TCR-T_MART-1_ were enriched with exhausted and anergic cells (Fig. 3E).

### Tumor PD-L1 expression resulted in various cellular and molecular responses in T cells

To correlate phenotypes other than cluster composition to T cell cytotoxicity (Fig. 1D), the expression of cytokines, chemokines, cytokine and chemokine receptors, and transcription factors was analyzed. With increased tumor PD-L1, expression of activation and cytotoxicity marker genes including *IFNG, TNFSF9, TNFSF14, CSF2*, and *IL2*, were downregulated in T_total_ (Fig. 4A), consistent with results of cytokine secretion assays (Fig. 1F, 1G). In addition, expression of anti-inflammatory cytokines, including *IL10, IL13*, and *IL19*, were upregulated in TCR-T_MART-1_ stimulated with PD-L1^high^ (Fig. 4A, S4A). Although the expression of pro-inflammatory cytokines such as *IL12A* and *IL5* was increased in TCR-T_MART-1_ targeting PD-L1^high^ (Fig. 4A), the results overall suggested the domination of anti-inflammatory cytokines over pro-inflammatory cytokines, resulting in the inhibition of T cell function. In line with the cytokine expression pattern, the expression of cytokine receptors related with T cell activation, including *TNFRSF9, IFNGR1*, and *IL2RA*, was upregulated in TCR-T_MART-1_ stimulated with PD-L1^low^ while the expression of *IL13RA2* and *IL13RA1* was increased in TCR-T_MART-1_ stimulated with PD-L1^high^ (Fig. 4B). Overall, the production of proinflammatory cytokines in TCR-T_MART-1_ was dose-dependently inhibited by the expression of PD-L1 in tumor cells and function of T cell targeting PD-L1^high^ was inhibited by the production of anti-inflammatory cytokines.

**Figure 4.**
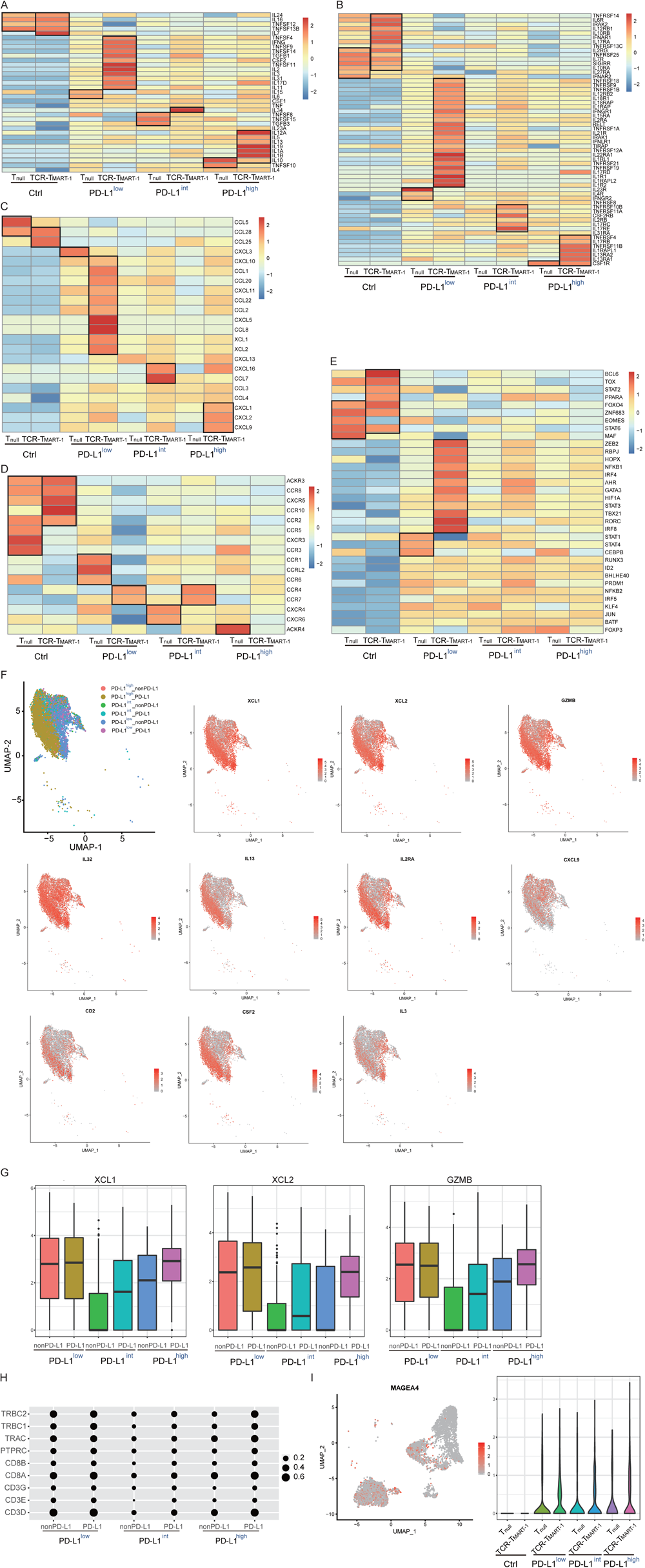
Gene expression of cytokine, chemokine, their receptors, and transcription regulators in T_null_ and TCR-T_MART-1_. **(A)** The expression file of cytokines in T_null_ and TCR-T_MART-1_. **(B)** The expression profile of cytokine receptors in T_null_ and TCR-T_MART-1_. **(C)** The expression file of chemokines in T_null_ and TCR-T_MART-1_. **(D)** The expression profile of chemokine receptors in T_null_ and TCR-T_MART-1_. **(E)** The expression profile of transcription factors in T_null_ and TCR-T_MART-1_. **(F)** UMAP projection of tumor cells, and the relative normalized expression of *XCL1, XCL2, GZMB, IL32, IL13, IL2RA, CXCL9, CD2, CSF2*, and *IL3*. **(G)** The boxplots showing the expression level of *XCL1, XCL2* and *GZMB* in nonPDL1 or PDL1-expressing tumor cells. **(H)** The bubble plot showing the expression of T cell marker genes in onPDL1 or PDL1-expressing tumor cells. **(I)** UMAP projection of T cells and the relative normalized expression of *MAGEA4* (left) and the violin plot showing the expression of *MEGEA4* in T_null_ and TCR-T_MART-1_ (right).

For chemokines (Fig. 4C), more were expressed in TCR-T_MART-1_ than in T_null_ after antigen stimulation. In Ctrl group, *CCL25* was highly expressed in TCR-T_MART-1_ but the expression of the CCL25 receptor gene *CCR9* was not detected. When cultured with PD-L1^low^, *CCL8* was highly expressed in TCR-T_MART-1_ (Fig. 4C) whereas *CCR2, CCR3, CCR5*, and, especially, *CCR1* that encoded CCL8 receptors were upregulated in T_null_ (Fig. 4D), indicating that TCR-T_MART-1_ could recruit T_null_ through chemokine secretion. As for PD-L1^int^, *CCL7* was significantly expressed in TCR-T_MART-1_, but *CCR1, CCR2*, and *CCR3* encoding CCL7 receptors were almost expressed equally low in both T_null_ and TCR-T_MART-1_. In PD-L1^high^, *CXCL2* and *CXCL9* were highly expressed in TCR-T_MART-1_, but *CXCR2* or *CXCR3* encoding their corresponding receptors were not detected or only weakly expressed (Fig. 4D). In conclusion, TCR-T_MART-1_ stimulated with PD-L1^low^ effectively attracted and activated T_null_, consistent with its greatest cytotoxicity.

Unique expression pattern of transcription factors (TFs) was also discovered in T_null_ and TCR-T_MART-1_ populations. The expression of *ZEB2, RBPJ, NFKB1, GATA3, IRF4* and *STAT3*, which are important for TCR signaling production and transduction and T cell activation and differentiation [36] [37], were higher in TCR-T_MART-1_ cultured with PD-L1^low^ (Fig. 4E). Overall, the expression of *IRF4, NFKB1*, and *RBPJ* was decreased in TCR-T_MART-1_ (Fig. S4B) whereas the expression of *EOMES* in T_null_ was progressively downregulated (Fig. S4C) with increasing percentages of PD-L1-expressing tumor cells. The results indicated the expression profile of transcription factors in T cells were also affected by tumor PD-L1 expression.

### Colocalization of tumor and TCR-T_MART-1_ increased immune cell cytotoxicity

Interestingly, some genes expressed by immune cells were detected in tumor cell clusters, including *XCL1, XCL2, GZMB, IL32, IL13, IL2RA, CXCL9, CD2, CSF2*, and *IL3* that are associated with T cell activation or cytotoxicity (Fig. 4F). This observation indicated that T cells and tumor cells were close enough or in contact when T cells were activated by tumor cells [38], thus were separated in the same droplet for scRNA-seq. To answer if this phenomenon accounted for the difference of T cell cytotoxicity caused by different percentages of PD-L1 expressing tumor cells, the above genes were assessed in tumor populations. Expression of cytotoxic genes *XCL1, XCL2*, and *GZMB* was highest in both T_null_ and TCR-T_MART-1_ cultured with PD-L1^low^ (Fig. 4G), in line with the highest cytotoxicity of T cells in PD-L1^low^ (Fig. 1D). Moreover, the expression of T cell marker genes including *CD3D, CD3E, CD3G, CD8A, CD8B, PTPRC, TRAC, TRBC1*, and *TRBC2* was also detected in tumor populations (Fig. 4H), confirming the presence of T cells in tumor populations.

The expression of the tumor cell marker gene *MAGEA4* was also detected in T cell populations, while no *MAGEA4* expression was detected in Ctrl group (Fig. 4I), suggesting the specificity of *MAGEA4* expression in tumor cells. In addition, the expression of *MAGEA4* was higher in TCR-T_MART-1_ than in T_null_ in each group and was the highest in TCR-T_MART-1_ cultured with PD-L1^low^, consistent with immune cell cytotoxicity (Fig. 1D).

### Increased expression of tumor PD-L1 enhanced T cell death

To detect the impact of PD-L1 expression on tumor and immune cell death, gene sets of cell death pathways, including apoptosis, necrosis, autophagy, pyroptosis, and ferroptosis, were used for GSVA analysis. We first analyzed tumor cells after they were cocultured with T cells for 24 h. Cell death pathways, especially necrosis and autophagy, were most enriched in PD-L1^int^ (Fig. 5A), suggesting a non-linear correlation between PD-L1 expression and tumor cell death at the transcriptional level. When tumor populations were seperated into PD-L1-expressing or PD-L1-non-expressing (nonPD-L1) subsets, cell death pathways were most enriched in PD-L1-expressing cells of PD-L1^int^ (Fig. 5B). The expression of the key members of these cell death pathways was further analyzed (Fig. 5C). Apoptotic genes, including *TRADD, BID, FAS, FASL*, autophagy gene *BECN1*, and ferroptosis genes *GLS2, VDAC3, CARS, GPX4, HSPB1, NFE2L2*, were upregulated in PD-L1-expressing tumor cells of PD-L1^low^ (Fig. 5C, S5A, S5B), providing a possible reason for the strongest cytotoxicity observed in PD-L1^low^ (Fig. 1D).

To further assess the difference between tumor populations of each group, GO analysis were performed in tumor cells. PD-L1-expressing populations in each group had similar enriched signaling pathways, including protein localization or targeting to endoplasmic reticulum (ER) and antigen processing and presentation pathways (Fig. 5D). In contrast, pathways enriched in nonPD-L1 populations varied from each other and from PD-L1-expressing subsets (Fig. 5D). To gain insight into whether immune cell death would be affected by tumor PD-L1, cell death pathways (Fig. 5E) and gene expression (Fig. S5C) were analyzed in T_null_ and TCR-T_MART-1_. Cell death pathways were more enriched in TCR-T_MART-1_ than in T_null_ in each group and the enrichment of cell death signaling was positively correlated with the level of tumor PD-L1 while negatively correlated with T cell cytotoxicity (Fig. 1D). These results suggested that tumor PD-L1 enhanced T_null_ and TCR-T_MART-1_ cell death, thus inhibited T cell function.

**Figure 5.**
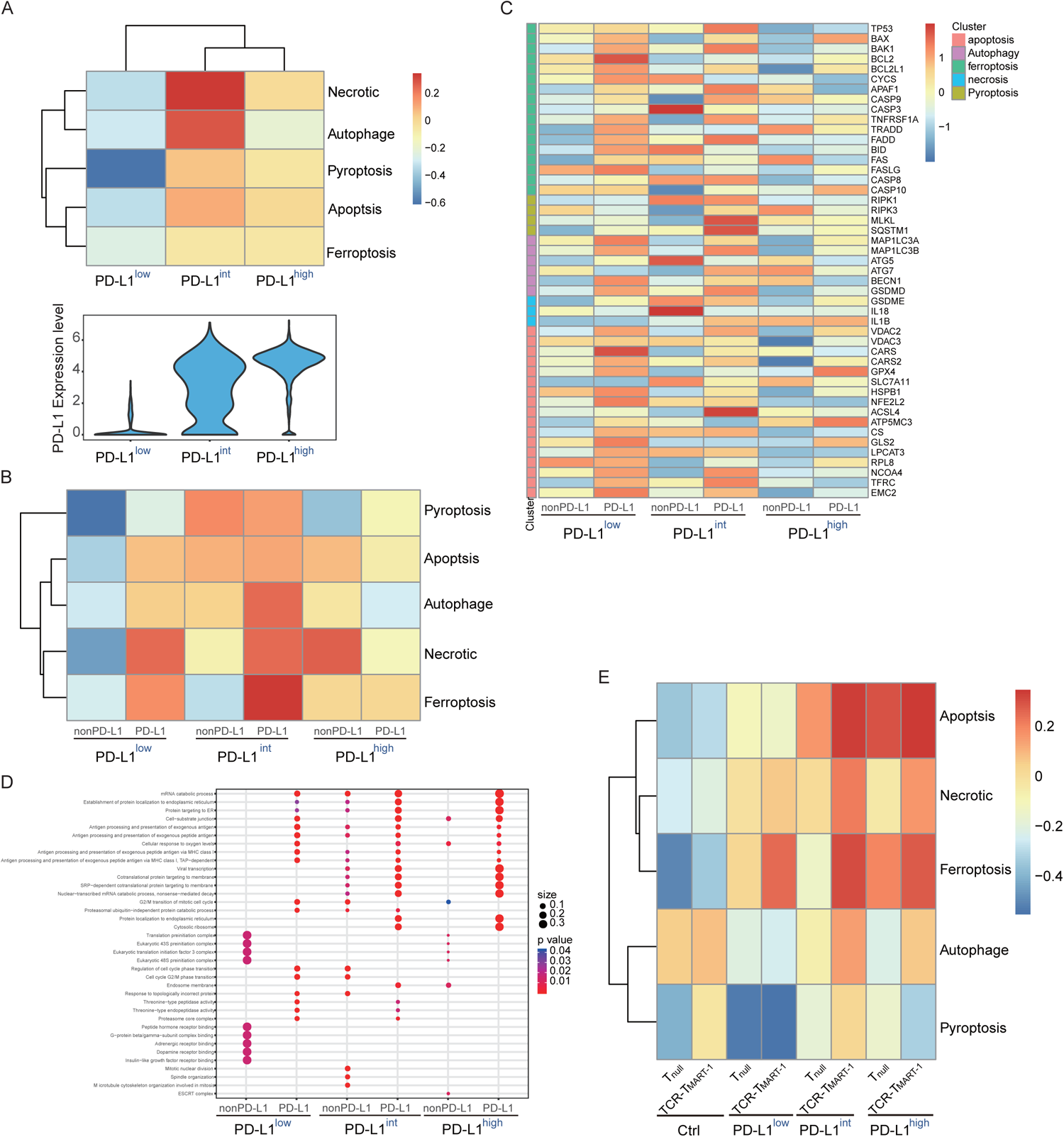
Enrichment of cell death pathways in tumor cells and T cells. **(A)** GSVA analysis of cell death pathways in tumor cells (top) and violin plot showing the expression level of PD-L1 in tumor cells (bottom). (**B**) GSVA analysis of cell death pathways in tumor cells expressing PD-L1 or not. (**C)** Heatmap of gene members from different cell death pathways. (**D)** Bubble plot showing the top 10 pathways enriched in different subsets of tumor cells. The color represents p value and the size represents gene ratio. (**E)** GSVA analysis of cell death pathways in different subsets of T cells.

### Tumor PD-L1 levels correlated positively with PD-L1 expression and negatively with PD-1 expression in T cells

PD-L1 has been reported to interact with CD80 in *cis* to selectively suppress the CD80:CTLA4 interaction but not the CD80:CD28 interaction [9]. To reveal how the PD-L1 network worked here, the expression of PD-L1, PD-1, CD80, CTLA4, and CD28 was assessed. With the increase of tumor PD-L1 (Fig. 6A, S6A), the proportion of PD-L1^+^ and the level of PD-L1 in T_null_ and, especially, in TCR-T_MART-1_ gradually increased (Fig. 6B, S6B). PD-1 expression was highest in TCR-T_MART-1_ targeting PD-L1^low^ (Fig. 6C, S6C) and decreased with increasing tumor PD-L1. It implied the strongest T cell activation induced highest PD-1 expression with lowest PD-L1 expression. Since CD80 expression in tumor cells (Fig. 6D) or T_total_ (Fig. 6E) was much lower than PD-L1 expression (Fig. 6A, 6B), CD80 would entirely bind to PD-L1 in *cis*, rather than to CTLA4 (Fig. 6F) in *trans*. PD-L1:CD80 *cis*-heterodimer could then trigger co-stimulatory receptor CD28 in T_null_ and TCR-T_MART-1_ (Fig. 6G). Considering the dramatic difference in expression levels of PD-L1 and CD80, the dominant signaling was the interaction between PD-L1 and PD-1 under the circumstances. Overall, the tumor PD-L1 level positively correlated with PD-L1 expression while negatively correlated with PD-1 expression on T cells, perfectly demonstrating that PD-1 expression is an activation marker for T cells [39].

**Figure 6.**
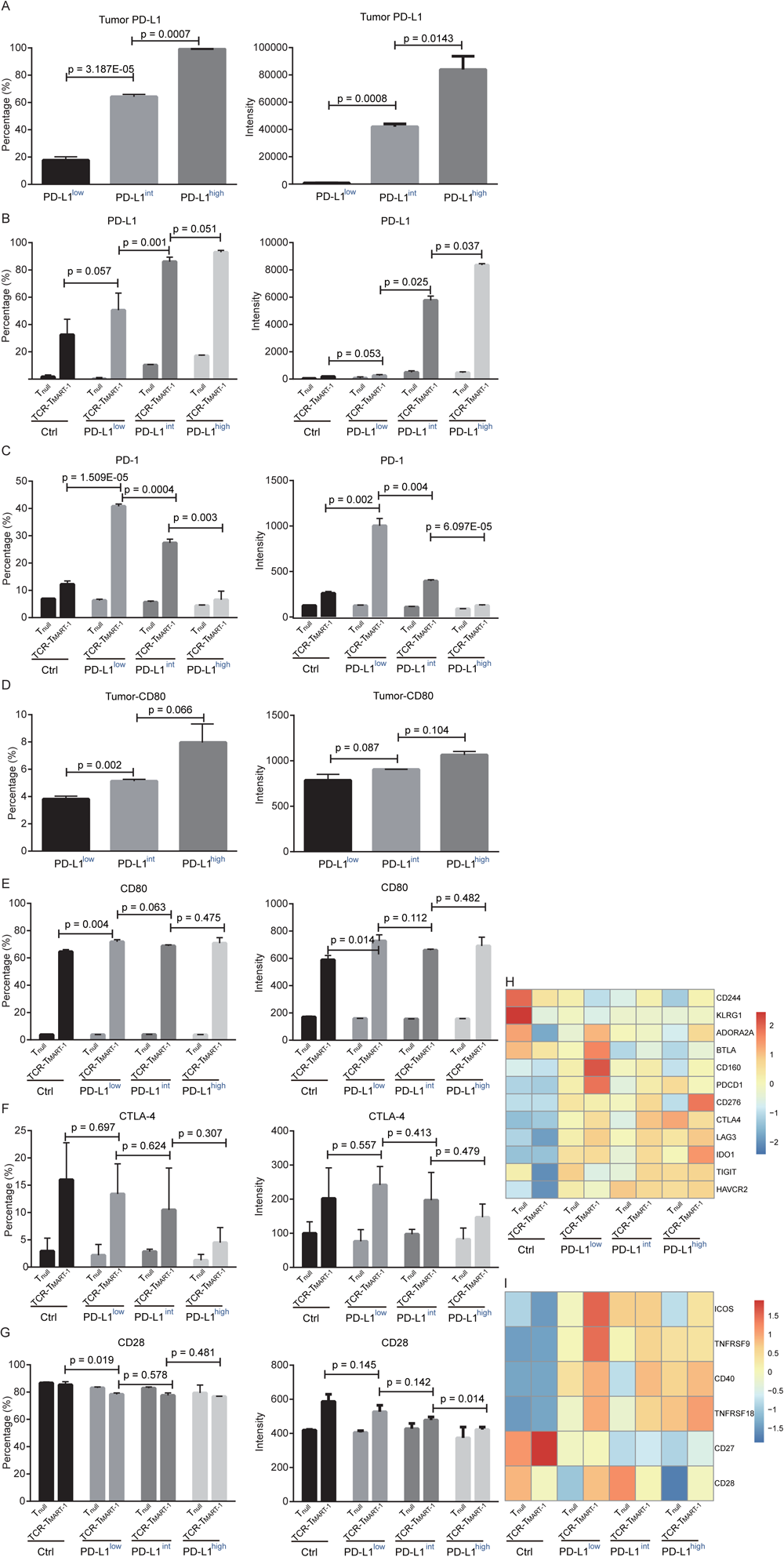
PD-L1 network and expression of immune checkpoint molecules in T cells. **(A)** The percentage (left) and intensity (right) of PD-L1 expression on T cells were assessed by FACS after incubation with MEL-526 cells for 24 h (n = 3). **(B)** The percentage (left) and intensity (right) of PD-L1 expression on tumor cells after incubation with MEL-526 cells for 24 h (n = 3). **(C)** The percentage (left) and intensity (right) of PD-1 expression on T cells after incubation with MEL-526 cells for 24 h (n = 3). **(D)** The percentage (left) and intensity (right) of CD80 expression on T cells after incubation with MEL-526 cells for 24 h (n = 3). **(E)** The percentage (left) and intensity (right) of CD80 expression on tumor cells after incubation with MEL-526 cells for 24 h (n = 3). **(F)** The percentage (left) and intensity (right) of CTLA-4 expression on T cells after incubation with MEL-526 cells for 24 h (n = 3). **(G)** The percentage (left) and intensity (right) of CD28 expression on T cells after incubation with MEL-526 cells for 24 h (n = 3). **(H)** Expression of inhibitory checkpoint molecules in T_null_ and TCR-T_MART-1_ with different ratios of PD-L1^+^ and PD-L1^-^ tumor cells. **(I)** Expression of stimulatory checkpoint molecules in T_null_ and TCR-T_MART-1_.

Blockade of PD-1 has been reported to lead to a compensatory upregulation of other checkpoint pathways[40], thus we analyzed whether increased tumor PD-L1 affected other checkpoint molecules. The expression of inhibitory checkpoint molecules (ICMs), including *ADORA2A, BTLA, CD160*, and *PDCD1*, was downregulated while the expression of *CD276* and *IDO1* was upregulated with the increased tumor PD-L1 (Fig. 6H). Simultaneously, the expression of stimulatory checkpoint molecules (SCMs) such as *ICOS* and *TNFRSF9* was highest in TCR-T_MART-1_ targeting PD-L1^low^ (Fig. 6I), consistent with its greatest cytotoxicity. Taking together, PD-L1 expression on tumor cells affected the expression of other inhibitory and stimulatory checkpoint molecules, which together impacted T cell function.

### Expression of *PDCD1* and *CD274* correlated with COVID-19 severity

The PD-1/PD-L1 signaling plays an essential role not only in regulating tumor immune responses but also in balancing homeostasis and tolerance in virus infection [41]. The current pandemic coronavirus disease 2019 (COVID-19) is caused by infection of severe acute respiratory syndrome coronavirus 2 (SARS-CoV-2) [42-44], where the role of PD-1/PD-L1 is currently unclear. Thus it is necessary to investigate how PD-1/PD-L1 signaling works during COVID-19 progress in order to deal with it. Publicly available data of bronchoalveolar cells from three moderate (M1-M3) and six severe (S1-S6) COVID-19 patients, and four healthy controls (HC1-HC4) were collected for analysis (66630 cells, Table S1) [45]. 31 clusters were identified by classical signature genes according to the reference (Fig. 7A) [45]. Expression of PD-1 and PD-L1 was first analyzed at the patient group level in different cell subpopulations, four trends of their expression dynamics were observed (Fig. 7B, 7C). *PDCD1* expression was gradually elevated in T cell, B cell, myeloid dendritic cells (mDCs), and macrophages from HC to mild cases then to severe patients. In the 2^nd^ trend, *PDCD1* expression was specifically increased in plasma cells and epithelial cells in severe patients but not in mild patients (Fig. 7B). For the 3^rd^ trend, *PDCD1* expression was upregulated in mild patients but slightly reduced in severe patients in NK and plasmacytoid dendritic cells (pDCs) (Fig. 7B). No expression of *PDCD1* was detected in mast cells and neutrophils in the 4^th^ trend (Fig. 7B). The *CD274* expression in macrophages, mast cells, pDC, and T cells (1^st^ trend) correlated well with COVID-19 severity and was specifically increased in plasma cells of severe patients (2^nd^ trend) (Fig. 7C). When analyzed at the individual level, expression of *PDCD1* and *CD274* was also elevated in mild and severe patients (Fig. S7A, S7B). Overall, *PDCD1* expression in T cells, B cells, mDCs, and macrophages and *CD274* expression in macrophages, mast cells, pDC, and T cells correlated well with COVID-19 severity. Furthermore, *PDCD1* and *CD274* expression was specifically increased in epithelial and plasma cells of severe patients. Inflammatory signaling participates in modulating PD-L1 expression, particularly, STAT1, which can be activated by IFNγ or interleukin 6 (IL-6), is a crucial regulator for PD-L1 expression [46, 47]. Furthermore, plasma IFNγ level [43] and the IL-6 level in bronchoalveolar lavage fluid (BALF) [45] were reported to be increased in COVID-19 patients. Consistently, *STAT1* was found upregulated in both mild and severe patients (Fig. 7D), suggesting increased *CD274* expression might at least partly resulting from increased *STAT1* level in COVID-19 patients.

**Figure 7.**
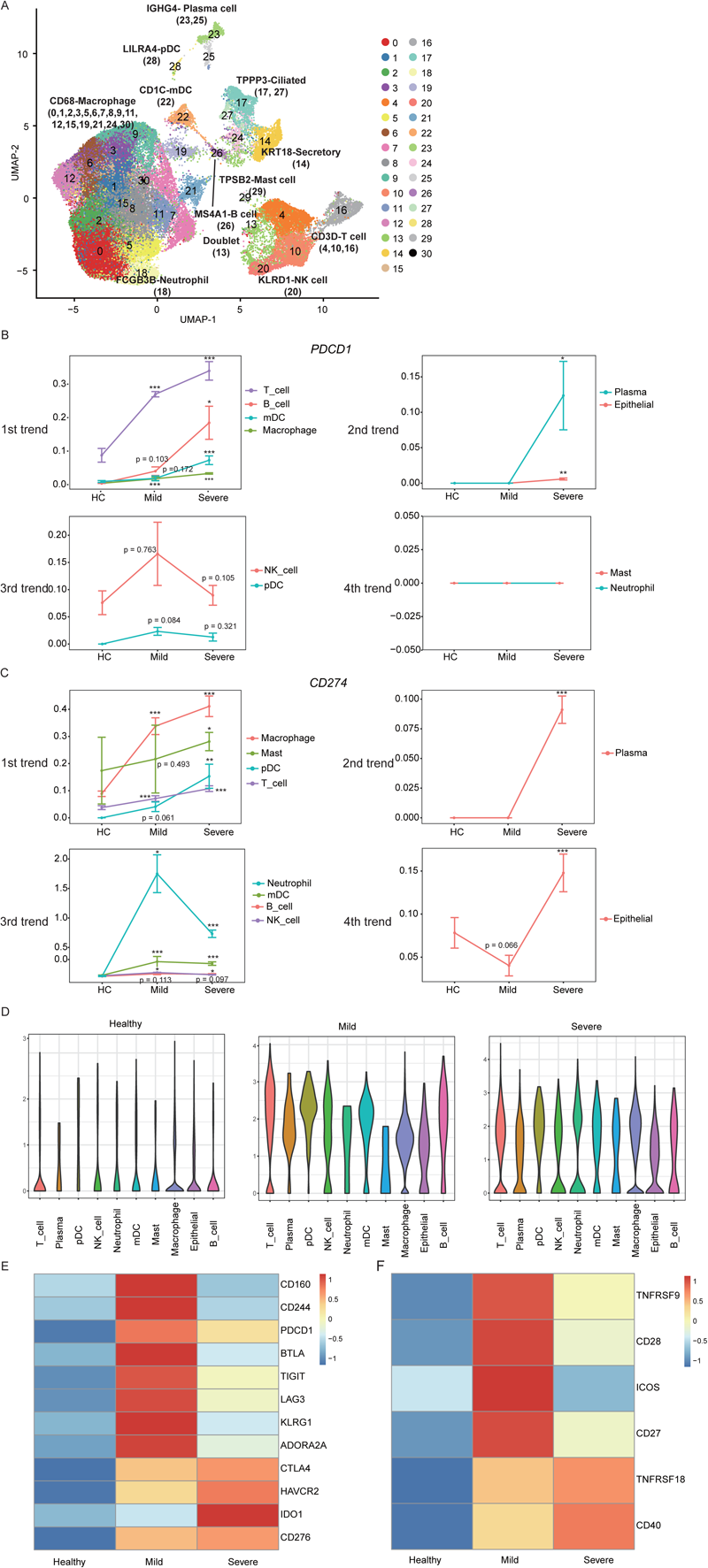
Single-cell immune profiling in COVID-19 patients. **(A)** The UMAP projection of BALF immune cells from HC and COVID-19 patients. **(B)** *PDCD1* expression in different cell subsets from HC and COVID-19 patients. **(C)** *CD274* expression in different cell subsets from HC and COVID-19 patients. **(D)** Violin plots showing the expression status of *STAT1* in different cell subsets from HC, Mild, and Severe COVID-19 patients. **(E)** Heatmap of inhibitory checkpoint molecules in HC as well as Mild and Severe COVID-19 patients. **(F)** Heatmap of stimulatory checkpoint molecules in HC as well as Mild and Severe COVID-19 patients.

To further elucidate the immune checkpoint landscape of COVID-19 patients, expression of classical inhibitory and stimulatory checkpoint molecules was assessed. For ICMs, expression of *CD160, CD244, PD-1, BTLA, TIGIT, LAG3, KLRG1*, and *ADORA2A* were increased in mild patients compared to HC and severe patients while expression of *CTLA4, HAVCR2, IDO1*, and *CD276* were highest in severe patients (Fig. 7E). Regarding SCMs, expression of *TNFRSF9, CD28, ICOS*, and *CD27* were elevated in mild patients in comparison to HC and severe patients while expression of *TNFRSF18* and *CD40* were highest in severe patients (Fig. 7F). When analyzed at the cell subpopulation level, unique expression patterns of ICMs and SCMs were demonstrated in each cell subpopulation (Fig. S7C, S7D).

## Discussion

It has been well documented that the efficacy of CPI in treating tumors is affected by the PD-L1 level, and the relative amounts of PD-1 and its interactors in tumors [9]. However, the molecular mechanism by which different levels of PD-L1 on tumors affect the therapeutic efficacy of TCR-T cell therapy remains unclear.

There are few studies about the effect of PD-L1 expression levels on TCR-T cell function. Our study provides an insight about TCR-T cell response to different proportions of tumor cells expressing PD-L1 at the single-cell level. The results of cell-based assays revealed that higher proportion of PD-L1^+^ tumor cells more strongly inhibited T-cell function (Fig. 1D-1G). Single-cell transcriptome profiling demonstrated the inhibition from different aspects, including cell differentiation (Fig. 3), secretion of cytokines and chemokines (Fig. 4A, 4C), colocalization of tumor and immune cells (Fig. 4F-4I), tumor and immune cell death (Fig. 5), as well as the whole PD-L1 network (Fig. 6A-6G).

TCR-T_MART-1_ were more vulnerable than T_null_ when targeting increasing proportion of PD-L1-bearing tumor cells (Fig. 3B-3C). It indicates that TCR-T therapy could be co-administrated with PD-L1/PD-1 interference to obtain better anti-tumor efficacy. Clinical trials with TCR-T cells armed with a PD-1 antagonist are ongoing (NCT04139057, NCT03578406). The result also implies that TCR-T cells will benefit from elimination of their *PDCD1*, such as by using CRISPR-based approaches, to protect themselves against PD-L1-mediated inhibition [48].

In addition to pro-inflammatory cytokines, the expression of anti-inflammatory cytokines, including *IL10*, was upregulated in TCR-T_MART-1_ targeting PD-L1^high^ (Fig. 4A, S4A). It was reported that IL-10 levels in serum and in ascites were increased after treatment with PD-1 inhibitor, implicating that IL-10 participated in undermining the efficacy of anti-PD-(L)1 therapies [49]. Thus, combined blockade of PD-L1 and IL10 may further enhance T-cell immunity [50, 51]. Interestingly, colocalization of T and tumor cells was detected and correlated negatively with the expression of tumor PD-L1 (Fig. 4F-4I). Colocalization of T and tumor cells was supported by the imaged interaction of T cells and APCs [38]. Therefore, tumor PD-L1 may inhibit T cell cytotoxicity through hindering the colocalization and interaction of antigen-specific T cells and tumor cells.

Various cell death pathways were involved in tumor and T cell death (Fig. 5, S5) and the enrichment of cell death signaling in T cells correlated positively with tumor PD-L1 expression level. This is consistent with a former study, where CD8^+^ T-cell apoptosis was promoted by PD-1 and PD-L1 upregulation [52], implying T cell death caused by PD-L1 signaling is one of the tumor evasion pathways.

Furthermore, the elevation of tumor PD-L1 expression dose-dependently increased the expression of PD-L1 in T cells (Fig. 6B, S6B), while PD-1 expression was dose-dependently decreased (Fig. 6C, S6C). PD-1 has been reported to induce apoptosis of antigen-specific T cells [53], but here tumor PD-L1 seems to play a dominant role in promoting effector T cell death. Moreover, consistent with that PD-1^+^ CD8^+^ T cells were functional cytotoxic T cells that targeted tumors and experienced exhaustion [54], PD-1 expression in T cells correlated positively with cytotoxicity in our study.

Lastly, since COVID-19 is pandemic and threatening thousands of people’s life, it is urgent and essential to investigate the molecular mechanism of the immune pathogenesis of the disease. Compared to healthy controls, *PDCD1* expression in T cells, B cells, mDCs, and macrophages (Fig. 7B) and *CD274* expression in macrophages, mast cells, pDCs, and T cells (Fig. 7C) were upregulated in COVID-19 patients, and correlated well with COVID-19 severity. Moreover, expression of *PDCD1* and *CD274* was specifically increased in plasma cells of severe patients (Fig. 7B, 7C), which could serve as a biomarker for prognosing the severity of COVID-19. Many clinical trials for treating COVID-19 are ongoing. Among them, one clinical trial uses PD-1 monoclonal antibody to block PD-1 in COVID-19 patients (NCT04268537). Based on our results, *PDCD1* expression was dramatically upregulated in T cells and macrophages especially in severe patients (Fig. 7B) and its blockade would further increase the secretion of multiple pro-inflammatory cytokines (Fig. 4A), which will enhance the cytokine release syndrome reported in COVID-19 patients and possibly associated with disease severity [42, 43], leading to further tissue damage or even more death especially in severe COVID-19 patients [55, 56]. A current study supports that checkpoint inhibitor immunotherapy is risky for severe outcomes in SARS-CoV-2-infected cancer patients, though these patients were treated with immune checkpoint inhibitors (ICI) before SARS-CoV-2 infection [57]. Furthermore, the expression of *IL10* was upregulated with increasing tumor PD-L1 (Fig. 4A), indicating a role of IL-10 in keeping a microenvironmental balance. Thus, the anti-inflammatory cytokine IL-10 might protect COVID-19 patients from T cell hyperactivation, which in turn may cause a dreaded complication characterized by acute respiratory distress syndromes in COVID-19 patients [58]. In addition, lower ratio of IL6 to IL10 serum levels was reported to coincided with the recovery of pneumonia [59].

In conclusion, cell-based cytotoxicity and cytokine secretion assays together with scRNA-seq were applied to interrogate MART-1-specific transgenic T cells upon antigen-specific stimulation with different ratios of tumor PD-L1. This study provides the first comprehensive illustration of tumor PD-L1 inhibition on TCR-T cell function at the single-cell level, and reveals some common mechanisms regarding how different subtypes of TCR-T cells respond to PD-L1 inhibition. It provides valuable information about the inhibition by increased tumor PD-L1 expression on TCR-T cells which are being applied in clinical trials, and about COVID-19, whose severity correlated well with the expression of *PDCD1* and *CD274*.

## Acknowledgments

We sincerely thank the support provided by China National GeneBank and Guangdong Provincial Key Laboratory of Genome Read and Write (No. 2017B030301011). This research was funded by Technology and Innovation Commission of Shenzhen Municipality, grant number JCYJ20170817150015170 and JSGG20180508152912700. This manuscript was edited by Life Science Editors.

## Author contributions

Q.G. and R.D. designed the project. Q.G. wrote the manuscript. Q.G. and C.C. revised the manuscript. S.L. performed the bioinformatic analysis. S.W., H.C. and Q.G. conducted the experiments related to single-cell sequencing. Q.X. and R.D. performed FACS analysis. F.W. and L.Z. assisted experiments and manuscript revision. R.D., H.C., and Q.G. performed all the other experiments and data analysis. Q.G., C.C., Y.G., and X.D. supervised the project.

## Methods

### Cell lines and cell culture

HEK293T (ATCC, CRL-11268) and T2 (174×CEM.T2, CRL-1992) cell lines were purchased from ATCC, and MEL-526 (BNCC340404) cell line was purchased from BNCC. HEK293T and MEL-526 cells were cultured in DMEM (Gibco, 21063029) supplemented with 10% fetal bovine serum (Hyclone, SH30084.03HI), penicillin (100 IU/mL), and streptomycin (50 μg/mL). T2 cells were cultured in IMDM (Gibco, 12440053) supplemented with 20% fetal bovine serum (Hyclone, SH30084.03HI), penicillin (100 IU/mL), and streptomycin (50 μg/mL). CD8^+^ T cells were cultured in HIPP-T009 (Bioengine, RG0101302) supplemented with 2% fetal bovine serum (Hyclone, SH30084.03HI) containing IL-2 (20 ng/ml), IL-7 (10 ng/ml) and IL-15 (10 ng/ml).

### Plasmid construction

TCR_MART-1_ sequence was identified from our previous work (data unpublished), and its constant regions were replaced by mouse TCR constant region α and β, respectively to prevent mispairing with endogenous TCR. TCR α chains and β chains were linked by P2A self-cleaving peptide. The recombinant DNA encoding TCR_MART-1_ was synthesized by GeneScript (Nanjing, China) and ligated into pRRLSIN.cPPT.PGK vector (Addgene, 12252).

PD-L1 cDNA ORF Clone in Cloning Vector was purchased from Sino Biological (HG10084). PD-L1 cDNA was cloned into pRRLSIN.cPPT.PGK vector (Addgene, 12252) with ClonExpress II One Step Cloning Kit (Vazyme, C112) according to the user manual.

### Lentivirus production

293T cells were transfected with a mixture of interested plasmid and packaging constructs (PsPAX2 and PMD2G) as previous [60]. The culture supernatants were collected 72 h after transfection and filtered through a 0.45 uM filter. Subsequently, the supernatants were concentrated by ultracentrifugation at 35,000 rpm for 90 min. The pellet was suspended and stored at -80°C.

### Generation of tumor cells expressing PD-L1

After lentivirus infection of PD-L1 lentivirus into MEL-526 cells for 2 days, PD-L1^+^ cells were sorted out by FACS. Different portions of PD-L1^+^ tumor cells were obtained by mixing wildtype and PD-L1^+^ MEL-526 cells.

### Generation of MART-1-specific T cells

Human Peripheral Blood Mononuclear Cells (PBMCs) were isolated from the blood of HLA-A*0201-restricted healthy donors with informed consent. CD8^+^ T cells were purified from PBMC via human CD8 MicroBeads (Miltenyi Biotec, 130-045-201) and activated with T Cell TransAct (Miltenyi Biotec, 130-111-160). After 36-48 h, CD8^+^ T cells were transduced with TCR_MART-1_ lentivirus at MOI=25 in a 6-well or 12-well plate. Simultaneously, polybrene was added to the culture at a final concentration of 2 μg/ml to promote infection efficiency. Then the well plate was centrifuge at 800g at room temperature for 30 minutes.

### Peptide synthesis

MART-1 originated peptide ELAGIGILTV (HLA-A*0201) was synthesized by GenScript (Nanjing, China) with a purity of ≥ 99.0%. Peptides were dissolved with 100% dimethyl sulfoxide (DMSO; Sigma-Aldrich, D5879-500ML) at the concentration of 10 mg/ml, and were stored at -20□.

### TCR-T cell stimulation with target tumor cell

TCR-T cells and MEL-526 cells (5×10^5 cells/ml concentration, in 200 μl) pulsed with peptide (final concentration 10 μg/mL) or not were incubated for 24 h in a round bottom 96-well plate. Afterwards, the co-culture was subjected to scRNA-seq. Unstimulated TCR-T cells (5×10^5 cells/ml) were incubated for 6 h alone before subjected to scRNA-seq.

### Intracellular staining

Cells were perforated and fixed using Cytofix/Cytoperm kit (BD Pharmingen, 554715).The antibodies used are as followed: Allophycocyanin(APC)-anti-HLA-A2 antibody (eBioscience, 17-9876-42), Phycoerythrin (PE)-anti-human CD8a antibody (eBioscience, 12-0086-42), APC-anti-human CD274(PD-L1) antibody (BD Pharmingen, 563741), PE-anti-human CD279(PD-1) antibody (Biolegend, 367404), PE anti-mouse TCR β chain Antibody (Biolegend, 109207), APC Anti-human IFN γ (eBioscience, 502512), PE-anti-human Granzyme B (BD Pharmingen, 561142), APC anti-human CD107a (Biolegend, 328620), PE-anti-Ki67 antibody (Abcam, ab270650).

### Cell killing assays

Target cells were labeled with Carboxyfluorescein succinimidyl ester (CFSE; Invitrogen) and co-cultured with 50% TCR-T cells at E:T ratio of 1:2. After 24 h, cells were collected and stained with PI and subsequently detected by FACS.

### Cytokine secretion measurement

The secretion of TNF-α, granzyme A, and granzyme B by T cell were evaluated using BDTM cytometric bead array (CBA) system. Tnull or TCR-T_MART-1_ cells were co-cultured with MEL-526 cells pulsed with peptide or not and supernatants were collected 24 h later. CBA assay was performed according to the instruction manual.

### Statistical analysis

Data analyzing was preformed using PRISM 6 (GraphPad Software) and RStudio. *P<0.05, **P<0.005, ***P<0.001. Values are presented as mean Standard deviation (SD). Error bars represented the SD.

### ScRNA-seq

Single-cell 3’ mRNA transcriptome profiling was performed using a negative pressure orchestrated DNBelab C4 system according to the workflow [31].

### ScRNA-seq data preprocessing

For all the samples, the iDrop Software Suite (v.1.0.0) was used to perform sample de-multiplexing, barcode processing and single-cell 3’ unique molecular identifier (UMI) counting with default parameters. Cleaned reads were then aligned onto the complete UCSC hg38 human genome by splicing-aware aligner STAR with default parameters. Valid cells were automatically identified based on the UMI number distribution of each cell. The filtering criteria were used to obtain high-quality single cell: the number of genes in each cell in the range of 400 to 6000, the ratio of mitochondrial genes less than 0.2, and the number of UMI more than 1000.

### Unsupervised clustering

The expression matrix obtained in the above steps was used as input to Seurat v. 3 to perform batch effect correction, standardization, dimensionality reduction, and clustering. First, the “LogNormalize” function was applied to normalize the data. Second, the “vst” method in the “FindVariableFeatures” function was used to detect variable genes, and the top 3000 variable genes were selected for downstream analysis. Third, the “FindIntegrationAnchors” and “IntegrateData” functions were used to correct batch effects. Fourth, the top 3000 variable genes were applied for PCA dimensionality reduction. The UMAP was performed on the top 20 principal components for visualizing these cells. At the same time, graph-based clustering was performed on the PCA-reduced data for clustering analysis with Seurat v.3. The resolution was set to 1 to obtain a most representative result.

### Differential gene expression analysis

We applied the FindMarkers to differential gene expression analysis. For each cluster of T cells and tumor cells, DEGs were generated relative to all of the other cells. A gene was considered significant with adjusted P< 0.05 and logFC > 0.25. To compare DEGs across CD8+ T cells and tumor cells under different experimental conditions, the limma method was used with the parameters recommended in the user guide for analysis. Then DEGs were identified when met these criteria: FDR adjusted p value of F test < 0.01.

### Developmental trajectory inference

The Monocle (version 2) algorithm with the signature genes of different functional clusters was applied to order CD8^+^ T cells excluding clusters expressing proliferating or mitochondrial genes in pseudo time. UMI value was first converted into normalized mRNA counts by the “relative2abs” function in monocle and created an object with parameter “expressionFamily = negbinomial.size” according to the Monocle tutorial. Then the CD8^+^ T cell differentiation trajectory was determined by the default parameters of Monocle.

### Gene set enrichment analysis

Gene Ontology (GO) enrichment analysis was performed on the differential genes of each cluster, and the results were used for cell type definition. The “enrichGO” function in the “clusterProfiler” package to perform GO analysis using the corresponding default parameters. Pathways with the q value <0.05 corrected by FDR were used for analysis.

### GSVA

GSVA was used to identify the molecular phenotype of each cluster with the normalized UMI data. The average normalized expression across T cell clusters was first obtained. Then, GSVA scores of gene sets for different clusters were calculated. GSVA values were plotted as a heatmap using R package “pheatmap”.

### Data availability

The data that support the findings of this study have been deposited into CNGB Sequence Archive (CNSA: https://db.cngb.org/cnsa/) of CNGBdb with accession number CNP0001109.

### Ethics approval and consent to participate

The study was approved by the Institutional Review Board on Bioethics and Biosafety of BGI. A written information consent was regularly obtained from all donors.

## Figure legend

**Figure S1. TCR**_**MART-1**_ **construction. (A)**. Schematic design of TCR_MART-1._ **(B)** Expression of TCR_MART-1_ on CD8^+^ T cells transfected by lentivirus before (middle) and after (right) cell sorting. **(C)** Killing of T2 cells by T_null_ and TCR-T_MART-1_ after co-incubation for 6 h at E:T ratio of 1:1. **(D)** PD-L1 was over expressed in MEL-526 cells. **(E)** Violin plot showing the expression of TCR_MART-1_ in different cell clusters.

**Figure S2. Volcano plot showing differentially expressed genes across T cell clusters**. Each red/blue dot denotes an individual upregulated/downregulated gene (logFC >= 1 and p.value < 0.01).

**Figure S3. Gene expression and signaling pathways in T cells responding to different expression of tumor PD-L1**. (**A)** The expression of DEGs in T_null_ targeting PD-L1^low^-, PD-L1^int^-, PD-L1^high^-expressing tumor cells (left) and the expression of DEGs in TCR-T_MART-1_ (right). **(B)** The bubble plot showing the top 5 pathways in TCR-T_MART-1_ targeting PDL1^low^-, PDL1^int^-, PDL1^high^ tumor cells.

**Figure S4. Expression of cytokines and transcription factors in T cells. (A)** The bar plot showing the average expression level of *IL10, IL13*, and *IL19* in T_null_ and TCR-T_MART-1_. **(D)** The bar plot showing the average expression level of *IRF4, NFKB1*, and *RBPJ* in T_null_ and TCR-T_MART-1_. **(E)** *EOMES* expression in T_null_ and TCR-T_MART-1_.

**Figure S5. Expression of cell death associated genes. (A)** The proportion of cells expressing *TRADD, BID, FAS, FASLG*, and *BECN1* in tumor cells. **(B)** The proportion of cells expressing *GLS2, VDAC3, CARS, GPX4, HSPB1*, and *NFE2L2* in tumor cells. (C) Heatmap showing the expression of cell death associated genes in T_null_ and TCR-T_MART-1_.

**Figure S6. Transcriptional profiles of *CD274* and *PDCD1* in T cells and tumor cells. (A)** Bar plot showing the average expression level (left) and percentage (right) of *CD274* in T _null_ and TCR-T_MART-1_. **(B)** Bar plot showing the average expression level (left) and percentage (right) of *CD274* in MEL-526 cells. **(C)** Bar plot showing the average expression level (left) and percentage (right) of *PDCD1* in T _null_ and TCR-T_MART-1_. **(D)** Bar plot showing the average expression level (left) and percentage (right) of *PDCD1* in MEL-526 cells.

**Figure S7. Immune profiling of checkpoint molecules in COVID-19 patients. (A)** *PDCD1* expression was upregulated in mild and severe COVID-19 patients compared to HC. Error bars represent ± standard error. **(B)** *CD274* expression was upregulated in mild and severe COVID-19 patients compared to HC. **(C)** Heatmap showing the expression pattern of inhibitory checkpoint molecules in different cell subsets from HC, mild and severe COVID-19 patients. **(D)** Heatmap showing the expression pattern of stimulatory checkpoint molecules in different cell subsets.

## References

1. W Zou, L Chen, Inhibitory B7-family molecules in the tumour microenvironment. Nat Rev Immunol, 2008. 8(6): p. 467–77.

2. Abiko K., Matsumura N., Hamanishi J., Horikawa N., Murakami R., Yamaguchi K., et al., IFN-gamma from lymphocytes induces PD-L1 expression and promotes progression of ovarian cancer. Br J Cancer, 2015. 112(9): p. 1501–9.

3. Mandai M., Hamanishi J., Abiko K., Matsumura N., Baba T., Konishi I., Dual Faces of IFNgamma in Cancer Progression: A Role of PD-L1 Induction in the Determination of Pro- and Antitumor Immunity. Clin Cancer Res, 2016. 22(10): p. 2329–34.

4. X Zhang, Y Zeng, Q Qu, J Zhu, Z Liu, W Ning, et al., PD-L1 induced by IFN-γ from tumor-associated macrophages via the JAK/STAT3 and PI3K/AKT signaling pathways promoted progression of lung cancer. International journal of clinical oncology, 2017. 22(6): p. 1026–1033.

5. H Dong, SE Strome, DR Salomao, H Tamura, F Hirano, DB Flies, et al., Tumor-associated B7-H1 promotes T-cell apoptosis: a potential mechanism of immune evasion. Nat Med, 2002. 8(8): p. 793–800.

6. Ghebeh H., Mohammed S., Al-Omair A., Qattan A., Lehe C., Al-Qudaihi G., et al., The B7-H1 (PD-L1) T lymphocyte-inhibitory molecule is expressed in breast cancer patients with infiltrating ductal carcinoma: correlation with important high-risk prognostic factors. Neoplasia, 2006. 8(3): p. 190–8.

7. Thompson R. H., Gillett M. D., Cheville J. C., Lohse C. M., Dong H., Webster W. S., et al., Costimulatory B7-H1 in renal cell carcinoma patients: Indicator of tumor aggressiveness and potential therapeutic target. Proc Natl Acad Sci U S A, 2004. 101(49): p. 17174–9.

8. Patsoukis N., Brown J., Petkova V., Liu F., Li L., Boussiotis V. A., Selective effects of PD-1 on Akt and Ras pathways regulate molecular components of the cell cycle and inhibit T cell proliferation. Sci Signal, 2012. 5(230): p. ra46.

9. Zhao Y., Lee C. K., Lin C. H., Gassen R. B., Xu X., Huang Z., et al., PD-L1:CD80 Cis-Heterodimer Triggers the Co-stimulatory Receptor CD28 While Repressing the Inhibitory PD-1 and CTLA-4 Pathways. Immunity, 2019. 51(6): p. 1059–1073 e9.

10. Mayoux M., Roller A., Pulko V., Sammicheli S., Chen S., Sum E., et al., Dendritic cells dictate responses to PD-L1 blockade cancer immunotherapy. Science Translational Medicine, 2020. 12: p. 1–11.

11. Porter D. L., Levine B. L., Kalos M., Bagg A., June C. H., Chimeric antigen receptor-modified T cells in chronic lymphoid leukemia. N Engl J Med, 2011. 365(8): p. 725–33.

12. Hamid O., Robert C., Daud A., Hodi F. S., Hwu W. J., Kefford R., et al., Safety and tumor responses with lambrolizumab (anti-PD-1) in melanoma. N Engl J Med, 2013. 369(2): p. 134–44.

13. Motzer R. J., Escudier B., McDermott D. F., George S., Hammers H. J., Srinivas S., et al., Nivolumab versus Everolimus in Advanced Renal-Cell Carcinoma. N Engl J Med, 2015. 373(19): p. 1803–13.

14. Robert C., Long G. V., Brady B., Dutriaux C., Maio M., Mortier L., et al., Nivolumab in previously untreated melanoma without BRAF mutation. N Engl J Med, 2015. 372(4): p. 320–30.

15. Borghaei H., Paz-Ares L., Horn L., Spigel D. R., Steins M., Ready N. E., et al., Nivolumab versus Docetaxel in Advanced Nonsquamous Non-Small-Cell Lung Cancer. N Engl J Med, 2015. 373(17): p. 1627–39.

16. Ferris Robert L., Blumenschein George, Fayette Jerome, Guigay Joel, Colevas A. Dimitrios, Licitra Lisa, et al., Nivolumab for Recurrent Squamous-Cell Carcinoma of the Head and Neck. New England Journal of Medicine, 2016. 375(19): p. 1856–1867.

17. Reck M., Rodriguez-Abreu D., Robinson A. G., Hui R., Csoszi T., Fulop A., et al., Pembrolizumab versus Chemotherapy for PD-L1-Positive Non-Small-Cell Lung Cancer. N Engl J Med, 2016. 375(19): p. 1823–1833.

18. Topalian S. L., S. Hodi, F., Brahmer J. R., Gettinger S. N., Smith D. C., McDermott D. F., et al., Safety, Activity, and Immune Correlates of Anti–PD-1 Antibody in Cancer. N Engl J Med, 2012. 366(26): p. 2443–54.

19. Herbst R. S., Soria J. C., Kowanetz M., Fine G. D., Hamid O., Gordon M. S., et al., Predictive correlates of response to the anti-PD-L1 antibody MPDL3280A in cancer patients. Nature, 2014. 515(7528): p. 563–7.

20. Daud Adil I., Wolchok Jedd D., Robert Caroline, Hwu Wen-Jen, Weber Jeffrey S., Ribas Antoni, et al., Programmed Death-Ligand 1 Expression and Response to the Anti–Programmed Death 1 Antibody Pembrolizumab in Melanoma. Journal of Clinical Oncology, 2016. 34(34): p. 4102–4109.

21. Larkin J., Chiarion-Sileni V., Gonzalez R., Grob J. J., Cowey C. L., Lao C. D., et al., Combined Nivolumab and Ipilimumab or Monotherapy in Untreated Melanoma. N Engl J Med, 2015. 373(1): p. 23–34.

22. Kluger H. M., Zito C. R., Turcu G., Baine M. K., Zhang H., Adeniran A., et al., PD-L1 Studies Across Tumor Types, Its Differential Expression and Predictive Value in Patients Treated with Immune Checkpoint Inhibitors. Clin Cancer Res, 2017. 23(15): p. 4270–4279.

23. Carlino Matteo S., Long Georgina V., Schadendorf Dirk, Robert Caroline, Ribas Antoni, Richtig Erika, et al., Outcomes by line of therapy and programmed death ligand 1 expression in patients with advanced melanoma treated with pembrolizumab or ipilimumab in KEYNOTE-006: A randomised clinical trial. European Journal of Cancer, 2018. 101: p. 236–243.

24. Tawbi H. A., Forsyth P. A., Algazi A., Hamid O., Hodi F. S., Moschos S. J., et al., Combined Nivolumab and Ipilimumab in Melanoma Metastatic to the Brain. N Engl J Med, 2018. 379(8): p. 722–730.

25. Garon E. B., Rizvi N. A., Hui R., Leighl N., Balmanoukian A. S., Eder J. P., et al., Pembrolizumab for the treatment of non-small-cell lung cancer. N Engl J Med, 2015. 372(21): p. 2018–28.

26. Hodi Frank Stephen, Chiarion-Sileni Vanna, Gonzalez Rene, Grob Jean-Jacques, Rutkowski Piotr, Cowey Charles Lance, et al., Nivolumab plus ipilimumab or nivolumab alone versus ipilimumab alone in advanced melanoma (CheckMate 067): 4-year outcomes of a multicentre, randomised, phase 3 trial. The Lancet Oncology, 2018. 19(11): p. 1480–1492.

27. Carbone D. P., Reck M., Paz-Ares L., Creelan B., Horn L., Steins M., et al., First-Line Nivolumab in Stage IV or Recurrent Non-Small-Cell Lung Cancer. N Engl J Med, 2017. 376(25): p. 2415–2426.

28. Butte M. J., Keir M. E., Phamduy T. B., Freeman G. J., Sharpe A. H., PD-L1 interacts specifically with B7-1 to inhibit T cell proliferation, in Immunity. 2007: Immunity. p. 111–122.

29. Li Y., Liu S., Hernandez J., Vence L., Hwu P., Radvanyi L., MART-1-specific melanoma tumor-infiltrating lymphocytes maintaining CD28 expression have improved survival and expansion capability following antigenic restimulation in vitro. J Immunol, 2010. 184(1): p. 452–65.

30. Yi M., Jiao D., Xu H., Liu Q., Zhao W., Han X., et al., Biomarkers for predicting efficacy of PD-1/PD-L1 inhibitors. Mol Cancer, 2018. 17(1): p. 129.

31. Liu Chuanyu, Wu Tao, Fan Fei, Liu Ya, Wu Liang, Junkin Michael, et al., A portable and cost-effective microfluidic system for massively parallel single-cell transcriptome profiling. bioRxiv preprint, 2019. doi: https://doi.org/10.1101/818450

32. Dufour J. H., Dziejman M., Liu M. T., Leung J. H., Lane T. E., Luster A. D., IFN-gamma-inducible protein 10 (IP-10; CXCL10)-deficient mice reveal a role for IP-10 in effector T cell generation and trafficking. J Immunol, 2002. 168(7): p. 3195–204.

33. Carr MW Roth SJ, Luther E, Rose SS, Springer TA, Monocyte chemoattractant protein 1 acts as a T-lymphocyte chemoattractant. Proc. Natl. Acad. Sci. USA, 1994. 91: p. 3652–56.

34. Trapnell C., Cacchiarelli D., Grimsby J., Pokharel P., Li S., Morse M., et al., The dynamics and regulators of cell fate decisions are revealed by pseudotemporal ordering of single cells. Nat Biotechnol, 2014. 32(4): p. 381–386.

35. Zheng C., Zheng L., Yoo J. K., Guo H., Zhang Y., Guo X., et al., Landscape of Infiltrating T Cells in Liver Cancer Revealed by Single-Cell Sequencing. Cell, 2017. 169(7): p. 1342–1356 e16.

36. Omilusik K. D., Best J. A., Yu B., Goossens S., Weidemann A., Nguyen J. V., et al., Transcriptional repressor ZEB2 promotes terminal differentiation of CD8+ effector and memory T cell populations during infection. J Exp Med, 2015. 212(12): p. 2027–39.

37. Dufva Olli, Koski Jan, Maliniemi Pilvi, Ianevski Aleksandr, Klievink Jay, Leitner Judith, et al., Integrated drug profiling and CRISPR screening identify essential pathways for CAR T-cell cytotoxicity. 2020. 135(9): p. 597–609.

38. Xiong W., Chen Y., Kang X., Chen Z., Zheng P., Hsu Y. H., et al., Immunological Synapse Predicts Effectiveness of Chimeric Antigen Receptor Cells. Mol Ther, 2018. 26(4): p. 963–975.

39. Simon S., Labarriere N., PD-1 expression on tumor-specific T cells: Friend or foe for immunotherapy? Oncoimmunology, 2017. 7(1): p. e1364828.

40. Huang R. Y., Francois A., McGray A. R., Miliotto A., Odunsi K., Compensatory upregulation of PD-1, LAG-3, and CTLA-4 limits the efficacy of single-agent checkpoint blockade in metastatic ovarian cancer. Oncoimmunology, 2017. 6(1): p. e1249561.

41. Qin Weiting, Hu Lipeng, Zhang Xueli, Jiang Shuheng, Li Jun, Zhang Zhigang, et al., The Diverse Function of PD-1/PD-L Pathway Beyond Cancer. Frontiers in Immunology, 2019. 10.

42. Chen Nanshan, Zhou Min, Dong Xuan, Qu Jieming, Gong Fengyun, Han Yang, et al., Epidemiological and clinical characteristics of 99 cases of 2019 novel coronavirus pneumonia in Wuhan, China: a descriptive study. The Lancet, 2020. 395(10223): p. 507–513.

43. Huang Chaolin, Wang Yeming, Li Xingwang, Ren Lili, Zhao Jianping, Hu Yi, et al., Clinical features of patients infected with 2019 novel coronavirus in Wuhan, China. The Lancet, 2020. 395(10223): p. 497–506.

44. Liu Y., Zhang C., Huang F., Yang Y., Wang F., Yuan J., et al., 2019-novel coronavirus (2019-nCoV) infections trigger an exaggerated cytokine response aggravating lung injury. ChinaXiv:202002.00018, 2020.

45. Liao M., Liu Y., Yuan J., Wen Y., Xu G., Zhao J., et al., Single-cell landscape of bronchoalveolar immune cells in patients with COVID-19. Nat Med, 2020.

46. Garcia-Diaz A., Shin D. S., Moreno B. H., Saco J., Escuin-Ordinas H., Rodriguez G. A., et al., Interferon Receptor Signaling Pathways Regulating PD-L1 and PD-L2 Expression. Cell Rep, 2017. 19(6): p. 1189–1201.

47. Moon J. W., Kong S. K., Kim B. S., Kim H. J., Lim H., Noh K., et al., IFNgamma induces PD-L1 overexpression by JAK2/STAT1/IRF-1 signaling in EBV-positive gastric carcinoma. Sci Rep, 2017. 7(1): p. 17810.

48. Gao Q., Dong X., Xu Q., Zhu L., Wang F., Hou Y., et al., Therapeutic potential of CRISPR/Cas9 gene editing in engineered T-cell therapy. Cancer Med, 2019. 8(9): p. 4254–4264.

49. Lamichhane P., Karyampudi L., Shreeder B., Krempski J., Bahr D., Daum J., et al., IL10 Release upon PD-1 Blockade Sustains Immunosuppression in Ovarian Cancer. Cancer Res, 2017. 77(23): p. 6667–6678.

50. Brooks D. G., Ha S. J., Elsaesser H., Sharpe A. H., Freeman G. J., Oldstone M. B., IL-10 and PD-L1 operate through distinct pathways to suppress T-cell activity during persistent viral infection. Proc Natl Acad Sci U S A, 2008. 105(51): p. 20428–33.

51. Sun Z., Fourcade J., Pagliano O., Chauvin J. M., Sander C., Kirkwood J. M., et al., IL10 and PD-1 Cooperate to Limit the Activity of Tumor-Specific CD8+ T Cells. Cancer Res, 2015. 75(8): p. 1635–44.

52. Shi F., Shi M., Zeng Z., Qi R. Z., Liu Z. W., Zhang J. Y., et al., PD-1 and PD-L1 upregulation promotes CD8(+) T-cell apoptosis and postoperative recurrence in hepatocellular carcinoma patients. Int J Cancer, 2011. 128(4): p. 887–96.

53. Francisco L. M., Sage P. T., Sharpe A. H., The PD-1 pathway in tolerance and autoimmunity. Immunol Rev, 2010. 236: p. 219–42.

54. Han J., Duan J., Bai H., Wang Y., Wan R., Wang X., et al., TCR Repertoire Diversity of Peripheral PD-1(+)CD8(+) T Cells Predicts Clinical Outcomes after Immunotherapy in Patients with Non-Small Cell Lung Cancer. Cancer Immunol Res, 2020. 8(1): p. 146–154.

55. Zhang C., Wu Z., Li J. W., Zhao H., Wang G. Q., Cytokine release syndrome in severe COVID-19: interleukin-6 receptor antagonist tocilizumab may be the key to reduce mortality. Int J Antimicrob Agents, 2020. 55(5): p. 105954.

56. Chua Robert Lorenz, Lukassen Soeren, Trump Saskia, Hennig Bianca P., Wendisch Daniel, Pott Fabian, et al., COVID-19 severity correlates with airway epithelium–immune cell interactions identified by single-cell analysis. Nat Biotechnol, 2020.

57. Robilotti E. V., Babady N. E., Mead P. A., Rolling T., Perez-Johnston R., Bernardes M., et al., Determinants of COVID-19 disease severity in patients with cancer. Nat Med, 2020.

58. Xu Z., Wang Y., Zhang J., Huang L., Zhang C., Liu S., et al., Pathological findings of COVID-19 associated with acute respiratory distress syndrome. Lancet Respir. Med., 2020. 8: p. 420–22.

59. de Brito R. C., Lucena-Silva N., Torres L. C., Luna C. F., Correia J. B., da Silva G. A., The balance between the serum levels of IL-6 and IL-10 cytokines discriminates mild and severe acute pneumonia. BMC Pulm Med, 2016. 16(1): p. 170.

60. Gao Q., Ouyang W., Kang B., Han X., Xiong Y., Ding R., et al., Selective targeting of the oncogenic KRAS G12S mutant allele by CRISPR/Cas9 induces efficient tumor regression. Theranostics, 2020. 10(11): p. 5137–5153.

