## Supplementary figures and images for "Single-cell transcriptome analysis of the immunosuppressive effect of differential expression of tumor PD-L1 on responding TCR-T cells"

### Supplementary figures

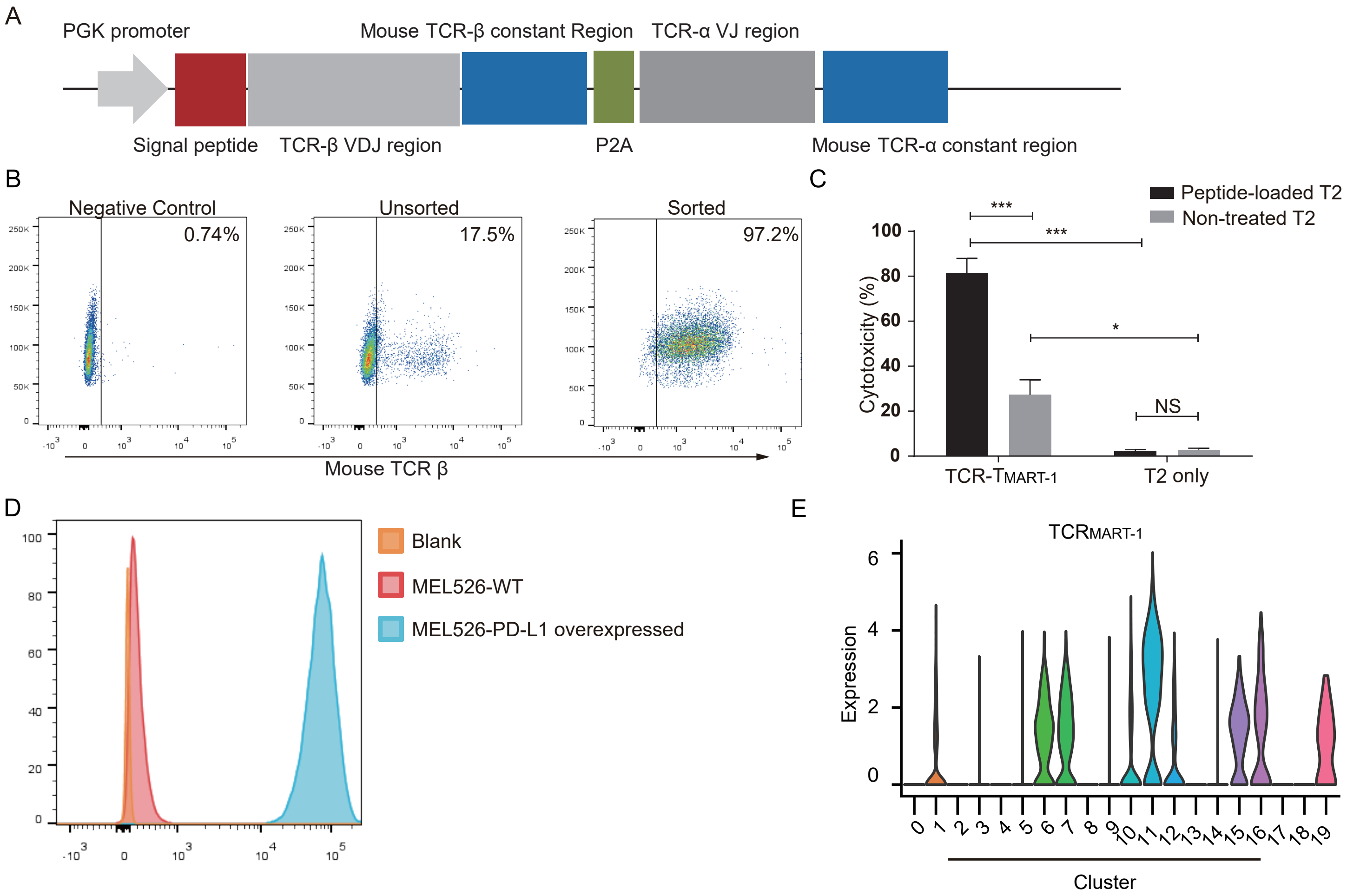

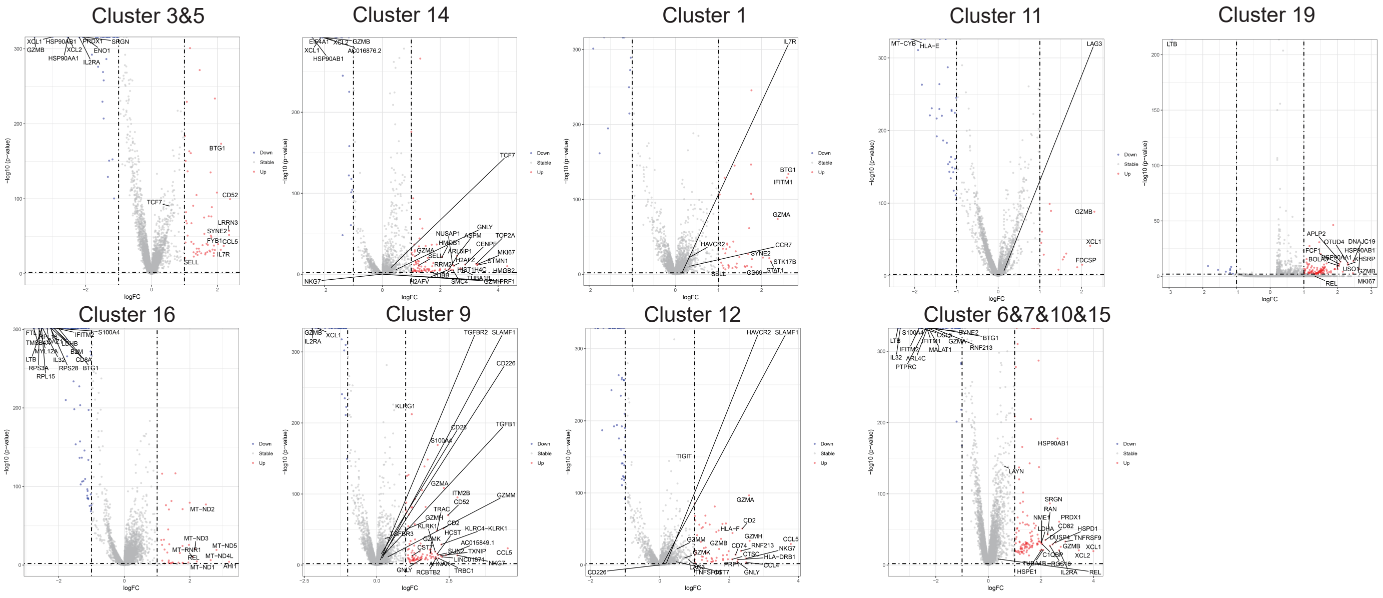

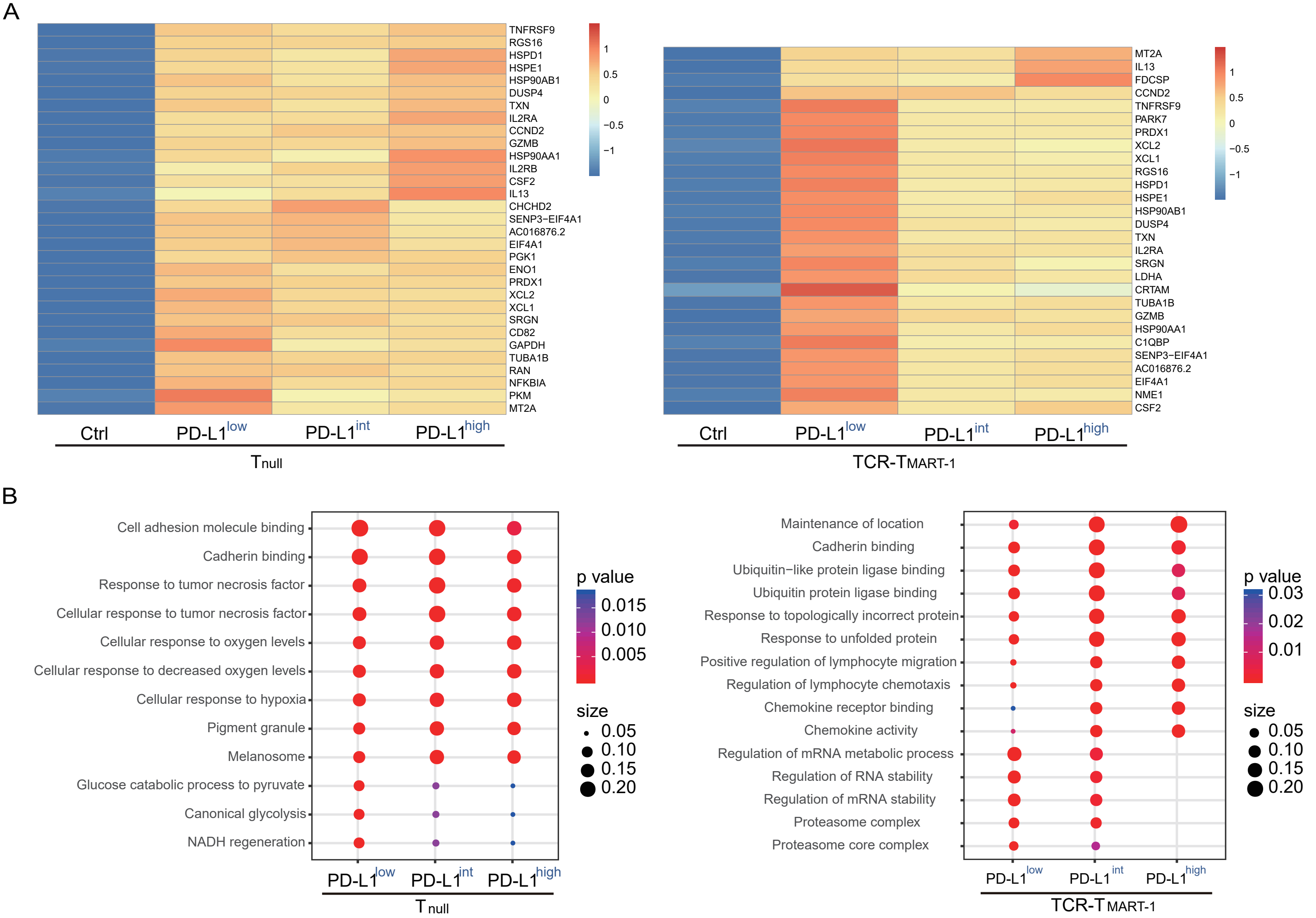

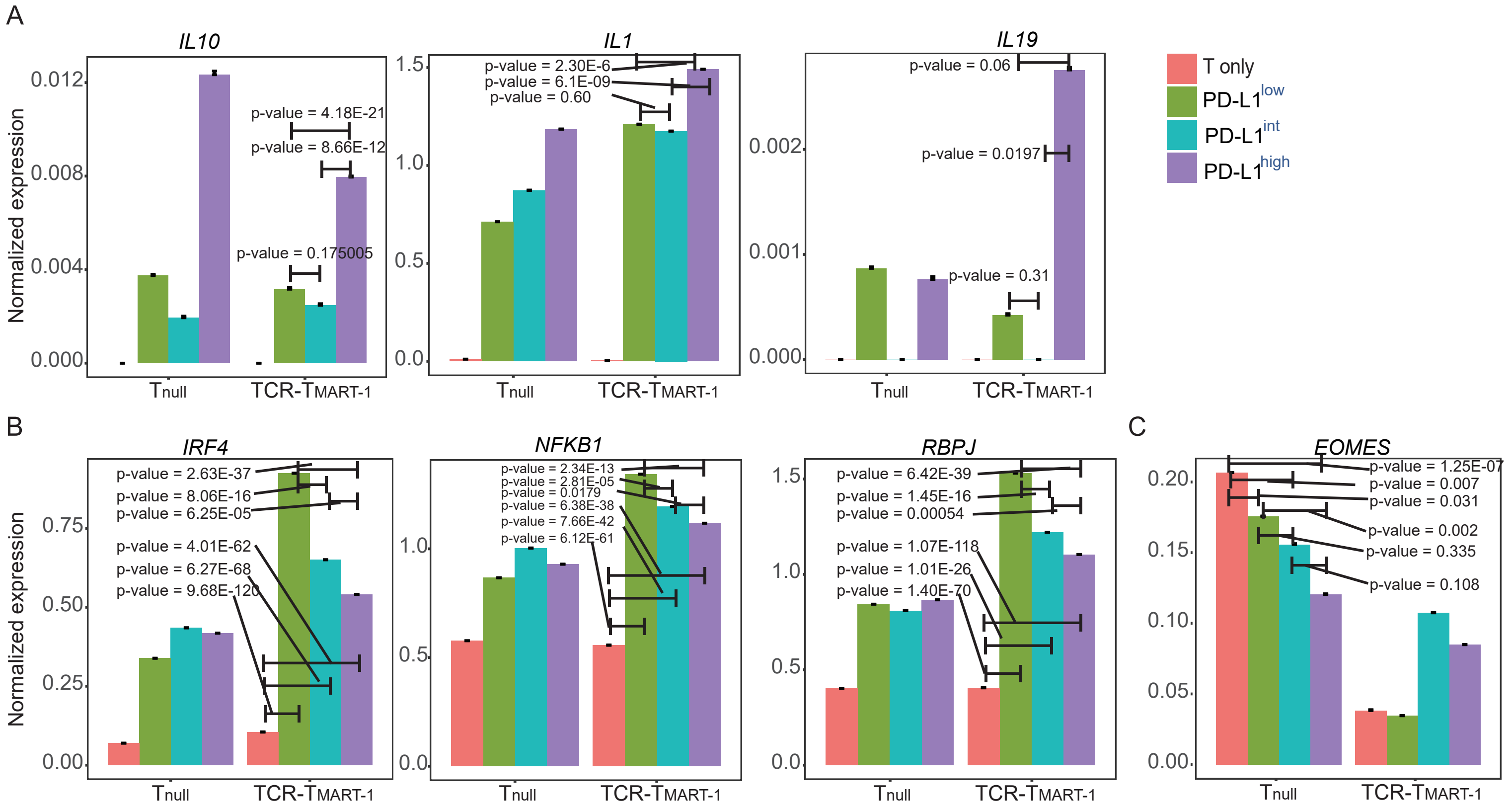

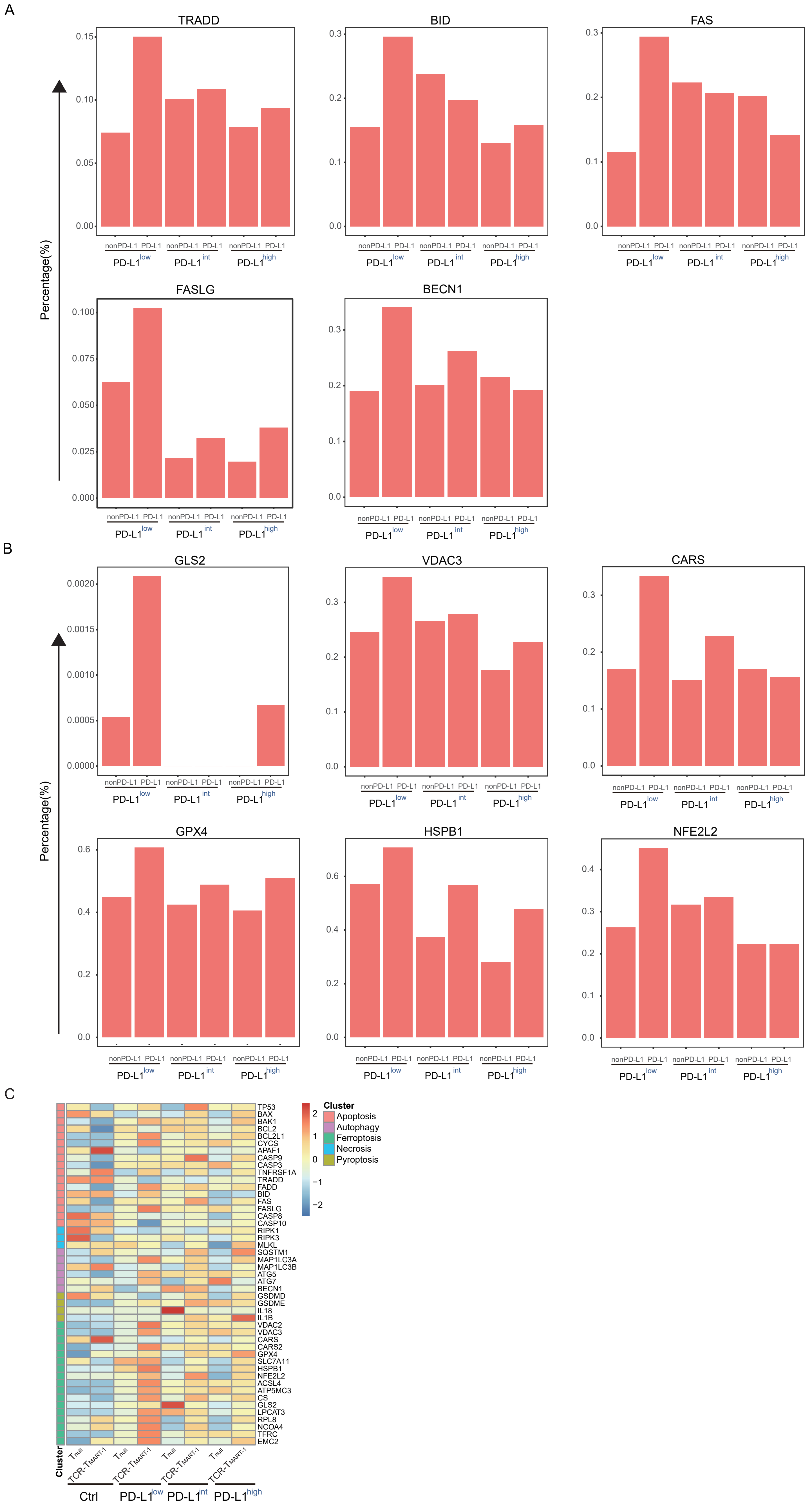

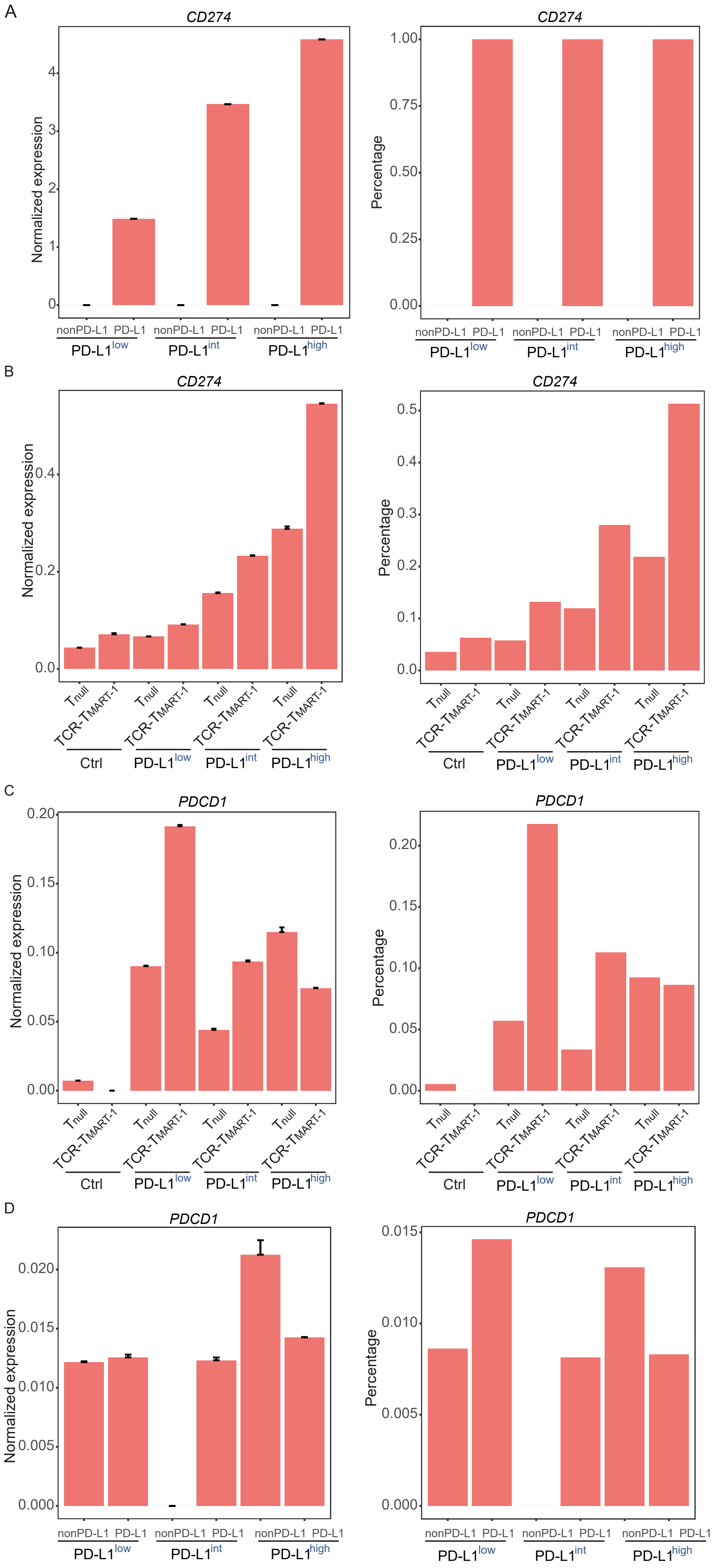

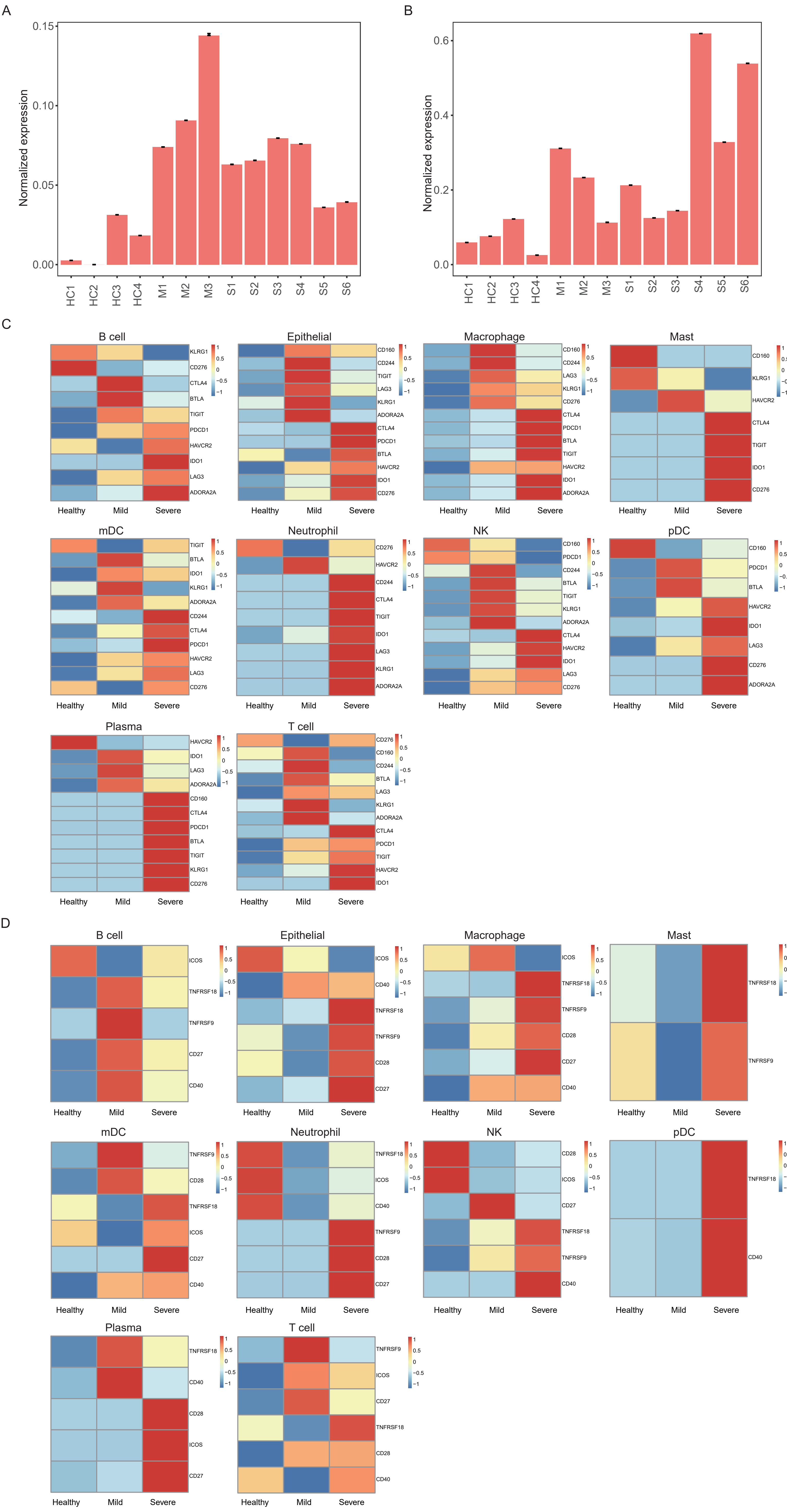

| Subject | Repeat analysis | In reference |
|---|---|---|
| C141 | 3542 | 3542 |
| C142 | 3411 | 3411 |
| C144 | 363 | 363 |
| C143 | 17340 | 17340 |
| C145 | 11872 | 11872 |
| C146 | 1292 | 1292 |
| C148 | 1718 | 1718 |
| C149 | 2071 | 2071 |
| C152 | 2904 | 2904 |
| C51 | 8644 | 8644 |
| C52 | 8189 | 8189 |
| C100 | 2566 | 2566 |
| GSM_3660650 | 2718 | 2718 |
| Total | 66630 | 66630 |
